# Evaluating crop models for future climate scenarios: wheat yield predictions using APSIM and STICS under combined CO_2_, warming, and water deficit conditions

**DOI:** 10.64898/2026.06.07.730737

**Authors:** Alan Severini, Meije Gawinowski, Marie-Odile Bancal, Marie Launay, Jean-Charles Deswarte, Karine Chenu

**Affiliations:** The University of Queensland, Queensland Alliance for Agriculture and Food Innovation (QAAFI), 5391 Warrego Hwy, Gatton, QLD 4343, Australia; Université Paris-Saclay, INRAE, AgroParisTech, UMR Ecosys, 91120 Palaiseau, France; INRAE, AgroClim, 84914 Avignon, France; Arvalis – Institut du végétal, Villiers-le-Bâcle, France; ARC Training Centre in Predictive Breeding, The University of Queensland, St Lucia, QLD 4072, Australia; ARC Hub for Engineering Plants to Replace Fossil Carbon, The University of Queensland, St Lucia, QLD 4072, Australia; ARC Centre of Excellence for Plant Success in Nature and Agriculture, The University of Queensland, St Lucia, QLD 4072, Australia

## Abstract

Crop models are essential for predicting climate change impacts on agriculture, yet their validation under multi-stress conditions remains limited. This study evaluated two widely-used wheat models, APSIM and STICS, using data from three Free-Air CO_2_ Enrichment (FACE) experiments (USA, Germany, Australia) combining elevated CO_2_ (eCO_2_), water deficit, and warming. Environmental characterisation using simulation-based stress indices revealed that intended “controls” frequently experienced hidden heat and water stress, meaning models were calibrated on crops already undergoing physiological adjustments. Evaluation of simulated yield and components revealed a clear hierarchy in prediction errors (RRMSE): unlimited conditions (3–9%) < single stress (4–27%, with a need to improve response to heat stress) < combined stress (17–123%). Elevated CO_2_ generally increased prediction uncertainty for crops experiencing water stress. Our results suggest that current stress functions from the models fail to capture the synergistic coupling between drought and heat stress. This highlights the urgent need for more mechanistic modelling to improve the reliability of climate change impact assessments.

## 1 Introduction

The concentration of carbon dioxide (CO_2_) in the atmosphere has increased in response to human activities, rising from 340 to 415ppm between 1980 and 2020. Future projections for the end of the century vary between 540 and 1300ppm (Lee et al., 2023). This increase in CO_2_ is accompanied by a rise in temperatures, which is predicted to result in higher average temperatures over the globe up to +4°C by 2100 (Calvin et al., 2023), and in an intensification of heat stress episodes. Extreme climatic events such as heatwaves (periods of abnormally high temperatures persisting over consecutive days) and droughts (prolonged periods of soil water deficit reducing crop water availability) are also set to become more frequent and intense. These climatic changes already affect crops with declining global yields for several species, such as wheat (Ababaei and Chenu, 2020; Ray et al., 2019). Moreover, further impacts are projected for the mid to long term (Collins and Chenu, 2021; Wang et al., 2023; Watson et al., 2017), with evidence that the frequency of the most stressful environment types has already increased in recent decades (Ababaei and Chenu, 2019; Collins and Chenu, 2022; Heidariask et al., 2026; Lake et al., 2025).

Crop response to isolated climatic factors such as water deficit (WD) and more recently increase in average temperature (‘high temperatures’, HT), heatwaves (HW) and elevated CO_2_ (eCO_2_) has been relatively thoroughly studied experimentally, yet not always for representative stress from production environments (Chenu, 2015; Lake et al., 2025; e.g. Rebetzke et al., 2012). By contrast, crop response to combined climatic factors remains understudied with persistent gaps in our understanding of stress interactions (Gawinowski et al., 2025b). Consistently, crop models have been reported to lack prediction capacity in those conditions (e.g. Chenu et al., 2017; Richetti et al., 2025). Experiments addressing these interactions are scarce. Even in commonly-studied wheat, only 26 studies have been reported for eCO_2_ × WD, 23 for eCO_2_ × HT, 9 for eCO_2_ × HW, 2 for eCO_2_ × HT × WD and 3 for eCO_2_ × HW × WD in a recent review on the subject (Gawinowski et al., 2025b). Moreover, these experiments were implemented for diverse genotypes and in very diverse conditions, including different ambient and elevated CO_2_ concentrations and stress characteristics (timing, intensity, duration), different types of facilities such as growth chambers, open-top chambers or FACE (Free-Air CO_2_ Enrichment). Due to these contrasted experimental conditions, crop responses widely vary between studies, highlighting the complexity of the interactions between the underlying physiological processes, and the need for proper robust modelling approaches (Gawinowski et al., 2025b).

Crop models are process-based models that are often associated with a soil model to simulate crop growth and development, soil water and nitrogen dynamics in response to environmental conditions and agricultural practices (Chenu et al., 2017). Since their development in the 1960s, these models have been widely used for diverse applications, including assessing and predicting the impacts of climate change on crops (Collins and Chenu, 2021; Lobell et al., 2015; Lobell and Asseng, 2017). Indeed, crop models are valuable tools for filling the data gaps from experimental studies and exploring the effects of climate variability on diverse processes. When robustly built on physiological processes and plant responses to the environment, such models can be used to extrapolate predictions to environmental scenarios beyond the range of current production systems (Chenu et al., 2017; Hammer et al., 2006).

Over recent decades, they have increasingly been used to make predictions under future climate scenarios (Chenu et al., 2017; Morel et al., 2021; Wang et al., 2023; Watson et al., 2017), although their prediction accuracy may vary depending on their framework and parameterisation. A critical challenge in using crop models to study the effects of climate change is their limited validation against experimental data, especially under combined fluctuations in CO_2_, temperature and water availability. In fact, only a very limited number of such validation has been reported, including validations for a few crop models (Ewert et al., 2002) and with crop model ensembles with AgMIP (Asseng et al., 2019) but only with a low range of environmental conditions tested. Given the high variation in mathematical framework and prediction outputs existing among crop models (Ahmed et al., 2019; Müller et al., 2021), individual crop model validation should be performed more systematically to allow robust predictions of climate change impacts, and appropriate assessment of adaptation strategies.

This study evaluates two crop models, APSIM and STICS, using data from previously-published field Free-Air CO_2_ Enrichment (FACE) experiments. The primary objective of this work was to assess the accuracy of these models in simulating wheat growth and development under varying environmental conditions including drought and heat stress in ambient and elevated atmospheric CO_2_. While an effort was made to incorporate diverse and comprehensive field-based datasets where different levels of CO_2_, soil water contents and thermal patterns were experienced, only a limited number of quality data with enough information for detailed model testing were identified. In their review, Gawinowski et al. (2025b) identified 63 experiments combining elevated CO_2_, warming and/or water deficit, conducted in different experimental facilities. Only 12 of these 63 experiments were conducted in FACE facilities, which are widely thought to best represent crop response in field conditions. Out of these 12 FACE experiments, data was only accessible for three which are the focus of the present study. These trials were conducted at Maricopa in the USA (Hunsaker et al., 2000; Kimball et al., 1999; Kimball et al., 1995), at Braunschweig in Germany (Manderscheid et al., 2020), and at Horsham in Australia (Fitzgerald et al., 2016). Tested treatments comprised a combination of CO_2_ levels ranging from 370 to 600ppm, rainfed and irrigated conditions, early and late sowing dates, and 0 to 3°C temperature increase during grain filling. For each location, the local cultivar grown in the trial was first parameterised in two crop models, APSIM and STICS, using data from the ‘control’ treatment (i.e. ambient CO_2_ level, no water stress, no increase in temperature, early sowing date). Models were then tested for the other studied conditions, which included elevated CO_2_, water stress and/or warmer conditions.

## 2 Materials and methods

### 2.1 Growing conditions and plant measurements

Three Free-Air Carbon Dioxide Enrichment (FACE) experiments on wheat (*Triticum aestivum* L.) with quality data were studied in this paper. They were performed at (i) Maricopa (USA) with spring wheat cultivar Yecora Rojo for a combination of CO_2_ × irrigation treatments (Kimball et al., 1995), (ii) Braunschweig (Germany) on winter wheat cv. Batis with a combination of CO_2_ × post-flowering heat treatments (Manderscheid et al., 2020), and Horsham (Australia) with spring wheat cv. Yitpi for a combination of CO_2_ × irrigation × sowing date treatments (Asseng et al., 2019; Fitzgerald et al., 2016; Mollah et al., 2009; O’Leary et al., 2015).

All experiments utilised the FACE technique to elevate atmospheric CO_2_ concentrations in open-field conditions, using rings of pipes that release CO_2_-enriched air around the crop canopy, with computer-controlled systems that adjust for wind speed and direction to maintain a stable target concentration across the plots.

It should be noted that maturity was explicitly defined as Zadoks stage 90 in the Horsham source data (Fitzgerald et al., 2016), whereas for Maricopa and Braunschweig only “maturity” was reported, without specifying whether this corresponded to physiological maturity or crop harvest; this inconsistency may introduce a small source of uncertainty in phenology comparisons across sites.

#### 2.1.1 Maricopa trials (different levels of CO_2_ and soil water patterns)

Two field experiments were conducted at the University of Arizona’s Maricopa Agricultural Center (MAC) in Maricopa, Arizona (33.5°N, 112.5°W) during the 1992-1993 and 1993-1994 growing seasons (Table 1). These trials were conducted to determine the combined effects of elevated atmospheric CO_2_ concentrations (eCO_2_) and water deficit (WD) on spring wheat *Triticum aestivum* L. cv Yecora Rojo (Kimball et al., 1995). The FACE setup maintained a target CO_2_ concentration of approximately 550ppm throughout the growing season for the elevated concentration, whereas the ambient CO_2_ concentration was at 370ppm. Two water treatments were imposed: an irrigated treatment, receiving water via a sub-surface drip irrigation system aimed at compensating for potential evapotranspiration, and a rainfed treatment, receiving only half the amount of water as the irrigated treatment at each irrigation event. To ensure that nutrients were not a limiting factor, all plots were fertilised with ample amounts of nutrients. Details of the trials and results can be found in Kimball et al. (1995).

**Table 1:** Characteristics of Maricopa trials and treatments, including the year, CO_2_ treatment, water regime, average flowering time (in days after sowing, ‘DAS’) and yield. The control treatments are presented in bold text.

| Year | CO<sub>2</sub> treatment | Water treatment | Flowering time (DAS) | Yield (g m<sup>-2</sup>) |
| --- | --- | --- | --- | --- |
| 1993 | Ambient (370ppm) | **Irrigated** | **102** | **804** |
| 1993 | Ambient (370ppm) | Rainfed | 101 | 677 |
| 1993 | Elevated (550ppm) | Rainfed | 98 | 735 |
| 1993 | Elevated (550ppm) | Irrigated | 100 | 917 |
| 1994 | Ambient (370ppm) | Rainfed | 117 | 619 |
| 1994 | Ambient (370ppm) | **Irrigated** | **121** | **768** |
| 1994 | Elevated (550ppm) | Rainfed | 116 | 742 |
| 1994 | Elevated (550ppm) | Irrigated | 117 | 860 |

#### 2.1.2 Braunschweig trials (different levels of CO_2_ and post-flowering temperatures)

Field experiments were conducted at the Johann Heinrich von Thünen-Institute, Federal Research Institute for Rural Areas, Forestry and Fisheries, Braunschweig, South-East Lower Saxony, Germany (52.30°N, 10.43°E) during the 2014 and 2015 growing seasons (Manderscheid et al., 2020). The study aimed to determine the combined effects of eCO_2_ and increased temperatures during the grain-filling period on winter wheat (*Triticum aestivum* L. cv Batis). The FACE setup imposed an elevated CO_2_ concentration targeted at 600ppm, whereas the ambient concentration was at 393ppm. Three temperature treatments were implemented with a T-FACE system with ceramic heaters: ambient conditions,moderate warming (+1.5°C), and high warming (+3°C) during grain filling, from 13 to 42 days after flowering in 2014, and from 5 to 37 days after flowering in 2015 (Table 2).

**Table 2:** Characteristics of Braunschweig trials and treatments, including the year, CO_2_ and temperature treatments, average flowering time (in days after sowing, ‘DAS’), and yield. The warming treatments were applied with ceramic heaters during the grain-filling period (‘GF’). The control treatments are presented in bold text.

| Year | CO<sub>2</sub> treatment | GF temperature increase (°C) | Flowering time (DAS) | Yield (g m<sup>-2</sup>) |
| --- | --- | --- | --- | --- |
| **2013** | **Ambient (393ppm)** | **0.0** | **220** | **785** |
| 2013 | Ambient (393ppm) | 1.5 | 220 | 795 |
| 2013 | Ambient (393ppm) | 3.0 | 220 | 733 |
| 2013 | Elevated (600ppm) | 0.0 | 220 | 867 |
| 2013 | Elevated (600ppm) | 1.5 | 220 | 937 |
| 2013 | Elevated (600ppm) | 3.0 | 220 | 818 |
| **2014** | **Ambient (393ppm)** | **0.0** | **219** | **785** |
| 2014 | Ambient (393ppm) | 1.5 | 219 | 722 |
| 2014 | Ambient (393ppm) | 3.0 | 219 | 783 |
| 2014 | Elevated (600ppm) | 0.0 | 219 | 879 |
| 2014 | Elevated (600ppm) | 1.5 | 219 | 921 |
| 2014 | Elevated (600ppm) | 3.0 | 219 | 844 |

Details of the trials and results can be found in Manderscheid et al. (2020).

#### 2.1.3 Horsham trials (different levels of CO_2_, soil water and thermal patterns)

The FACE field experiments at Horsham, Australia (36.8°S, 142.1°E, 128 m elevation) were conducted from 2007 to 2009. The experiments investigated the effects of two atmospheric CO_2_ concentrations (ambient at 385ppm and elevated at 550ppm), two water regimes (rainfed and supplemental irrigation), and two sowing times (early and late) to create different temperature environments during the growing season (Table 3) with the spring wheat cultivar Yitpi (O’Leary et al., 2015). Details of the trials can be found in Mollah et al. (2009), O’Leary et al. (2015), Fitzgerald et al. (2016), and Asseng et al. (2019). Data was obtained from Tables 1 and S1 in Fitzgerald et al. (2016).

**Table 3:** Characteristics of the Horsham trials and treatments, including the year, CO_2_, water and sowing date treatments, average flowering time (in days after sowing, ‘DAS’), and yield. Flowering time was only available for the treatment combination of ambient CO_2_, supplemental irrigation for both early and late sowings. The control treatments are presented in bold text.

| Year | CO<sub>2</sub> treatment | Water treatment | Sowing | Flowering time (DAS) | Yield (g m<sup>-2</sup>) |
| --- | --- | --- | --- | --- | --- |
| **2007** | **Ambient (385ppm)** | **Irrigated** | **Early (18 June)** | **133** | **359** |
| 2007 | Ambient (385ppm) | Irrigated | Late (23 Aug) | 89 | 237 |
| 2007 | Ambient (385ppm) | Rainfed | Late (23 Aug) | – | 206 |
| 2007 | Elevated (550ppm) | Irrigated | Early (18 June) | – | 406 |
| 2007 | Elevated (550ppm) | Irrigated | Late (23 Aug) | – | 303 |
| 2007 | Elevated (550ppm) | Rainfed | Late (23 Aug) | – | 226 |
| **2008** | **Ambient (385ppm)** | **Irrigated** | **Early (4 June)** | **138** | **357** |
| 2008 | Ambient (385ppm) | Irrigated | Late (5 Aug) | 93 | 170 |
| 2008 | Ambient (385ppm) | Rainfed | Late (5 Aug) | – | 167 |
| 2008 | Elevated (550ppm) | Irrigated | Early (4 June) | – | 495 |
| 2008 | Elevated (550ppm) | Irrigated | Late (5 Aug) | – | 180 |
| 2008 | Elevated (550ppm) | Rainfed | Late (5 Aug) | – | 149 |
| **2009** | **Ambient (385ppm)** | **Irrigated** | **Early (23 June)** | **126** | **242** |
| 2009 | Ambient (385ppm) | Irrigated | Late (19 Aug) | 86 | 133 |
| 2009 | Ambient (385ppm) | Rainfed | Late (19 Aug) | – | 111 |
| 2009 | Elevated (550ppm) | Irrigated | Early (23 June) | – | 358 |
| 2009 | Elevated (550ppm) | Irrigated | Late (19 Aug) | – | 229 |
| 2009 | Elevated (550ppm) | Rainfed | Late (19 Aug) | – | 158 |

### 2.2 Crop-model simulations

#### 2.2.1 APSIM and STICS

Simulations were conducted with APSIM (commit 820e02734, master branch, https://github.com/APSIMInitiative/ApsimX; Holzworth et al. (2018)) and version V10.1.0 of STICS (Beaudoin et al., 2023). The main growth processes affected by CO_2_, temperature (associated with heat) and water deficit are summarised in Table 4. The mathematical equations for the temperature response functions and the APSIM water-deficit senescence function are provided in Supplementary Note S1.

**Table 4:**
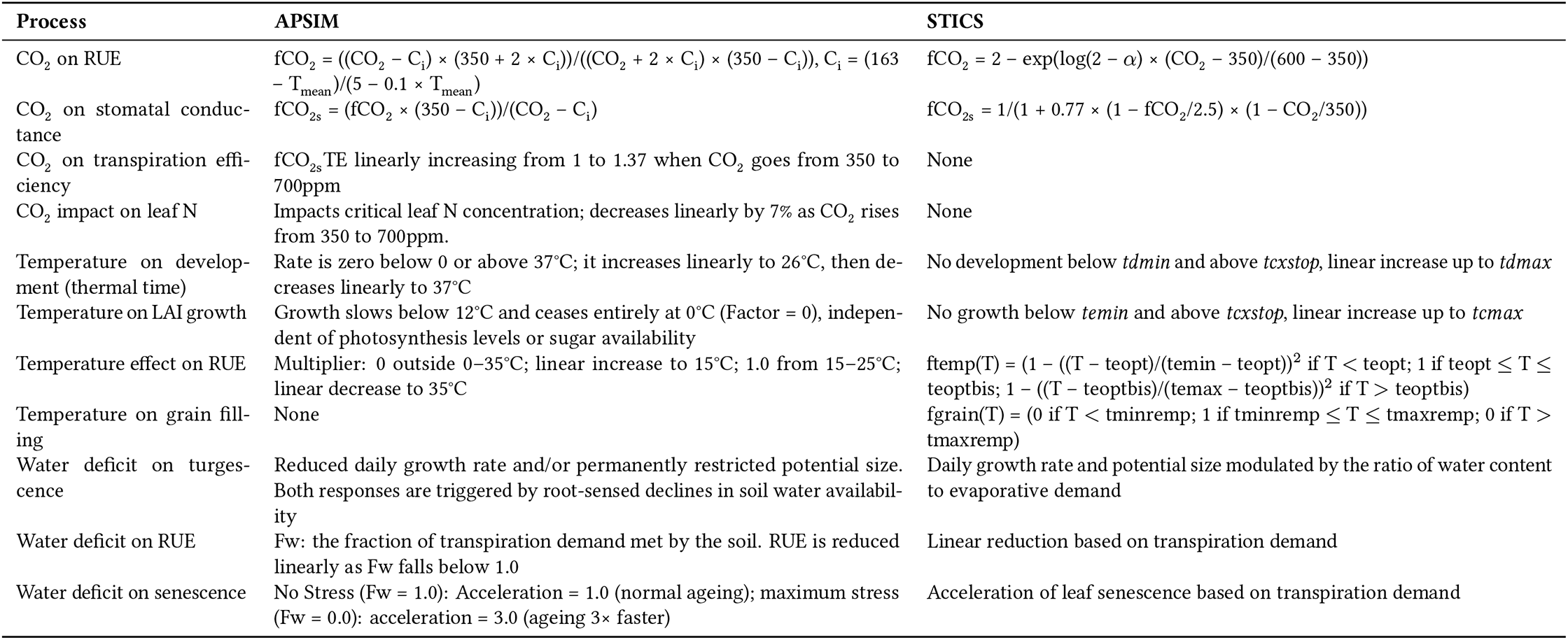
Summary of the main CO_2_, temperature (excluding low temperature and frost impacts) and water deficit direct effects on plant processes for APSIM and STICS. Plant parameters are represented in italic.

Evapotranspiration was also calculated differently between the two models. In APSIM, potential evapotranspiration and radiation partitioning are governed by the MicroClimate model, which uses a Penman-Monteith approach and a resistive network to calculate energy balances. For a standard wheat simulation, the canopy is represented as a single layer (from ground level to plant height), with soil evaporation calculated separately at the surface boundary; additional layers are only created when the physical structure of the canopy requires vertical discretisation (e.g., intercropping). Soil evaporation follows a Ritchie-style two-stage model, transitioning from a constant-rate phase to a falling-rate phase limited by hydraulic properties (so-called U and ConA coefficients), a method conceptually identical to the two-phase approach in STICS. In the default option of APSIM, root water uptake is handled by a Soil Arbitrator using a layer-by-layer extraction method based on ‘KL’ efficiency factors (https://www.apsim.info/apsim-wheat-documentation/; Zheng et al. (2015)).

In STICS, potential evapotranspiration is based on the resistive model of Shuttleworth and Wallace (Brisson et al., 1998) and the partitioning between transpiration and evaporation follows Beer’s law, where potential soil evaporation is attenuated by the canopy as *E*_*p,soil*_ = *E*_*p*_ ×*e*^*-k*×*LAI*^, with *k* an extinction coefficient. Potential soil water evaporation is based on the two phases model of Brisson et al. (1998), and root water uptake is simulated water absorption in the root zone distributed cm per cm according to the effective root profile and the available water content.

#### 2.2.2 Trial simulations and environmental characterisation

To replicate in silico the three experiments described in the previous subsection, trial management practices, soil characteristics and climatic data were retrieved from the respective source publications. Where weather data was incomplete, we supplemented with data obtained from NASA Power using the nasapower R package (Sparks, 2018).

Drought patterns were characterized using the commonly used water stress index (WSI) derived from APSIM simulations (Amin et al., 2025; e.g. Chapman et al., 2000; Chenu et al., 2013; Chenu et al., 2011; Christopher et al., 2016; Collins et al., 2021). This index corresponds to the soil “water supply-to-demand ratio” and indicates the extent to which soil water extractable by the roots (“water supply”) meets potential crop transpiration (“water demand”). Water demand represents the amount of water the crop would transpire in the absence of soil water limitation and is estimated daily as a function of crop growth and atmospheric vapour pressure deficit (Chenu et al., 2013). A WSI value of 1 indicates no water limitation, whereas a value of 0 indicates no water availability to the crop.

Heat stress events were characterized for the pre- and post-flowering periods identified as most impactful for wheat (Ullah et al., 2024). Specifically, heat stress was quantified as (i) the number of days with maximum temperature reaching 28 °C during the critical pre-flowering period between 300 to 200 °Cd before flowering, and (ii) the number of days with maximum temperature reaching 32 °C during early-to-mid grain filling, from 0 to 500 °Cd after flowering (Ullah et al., 2024; Yahya et al., 2026).

#### 2.2.3 Model calibration and validation

Studied cultivars were parameterised for APSIM and STICS using data from the control treatments of each trial. Key cultivar parameters were adjusted using observations for phenology, LAI, biomass, thousand grain weight and yield. STICS and APSIM parameters are presented in Supplementary Table S1 as well as in Supplementary Figure S1.

The accuracy of crop model simulations was assessed under eCO_2_, higher temperatures and/or limited irrigation treatments through correlations as well as the computation of root mean square errors (RMSE) and relative RMSE (RRMSE in %, calculated as the RMSE divided by the mean observed value and multiplied by 100) for different variables *v*, namely days to flowering, days to maturity, LAI, plant biomass, grain number and grain yield for each condition *i* and time (day) *t*. It should be noted that maturity was explicitly defined as Zadoks stage 90 in the Horsham source data (Fitzgerald et al., 2016), whereas for Maricopa and Braunschweig only “maturity” was reported, without specifying whether this corresponded to physiological maturity or crop harvest; this inconsistency may introduce a small source of uncertainty in phenology comparisons across sites.

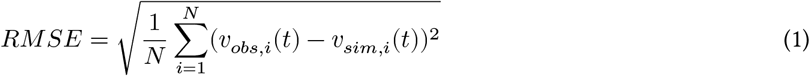

where *v*_*obs*_ is the observed value of the variable *v, v*_*sim*_ is its simulated value, and *N* is the number of observed values for *v*.

## 3 Results

### 3.1 Contrasting heat, drought and CO_2_ conditions

Environmental characterisation across the experimental sites revealed significant differences in heat and drought stress exposure during critical developmental phases. In particular, daily maximum temperature dynamics revealed a surprising contrast between Maricopa and Braunschweig regarding the actual heat stress experienced during grain filling, which did not align with the expected treatments.

In Braunschweig, where heat stress was intended as a treatment, maximum temperatures during the critical grain filling period barely exceeded 30°C (Fig. 1). Even with the experimental temperature increase of +3.0°C during grain filling, only 2 days during grain filling exceeded 32°C in both 2013 and 2014, while almost no stressful temperature was recorded pre-flowering (1 day; Fig. 2).

**Figure 1:**
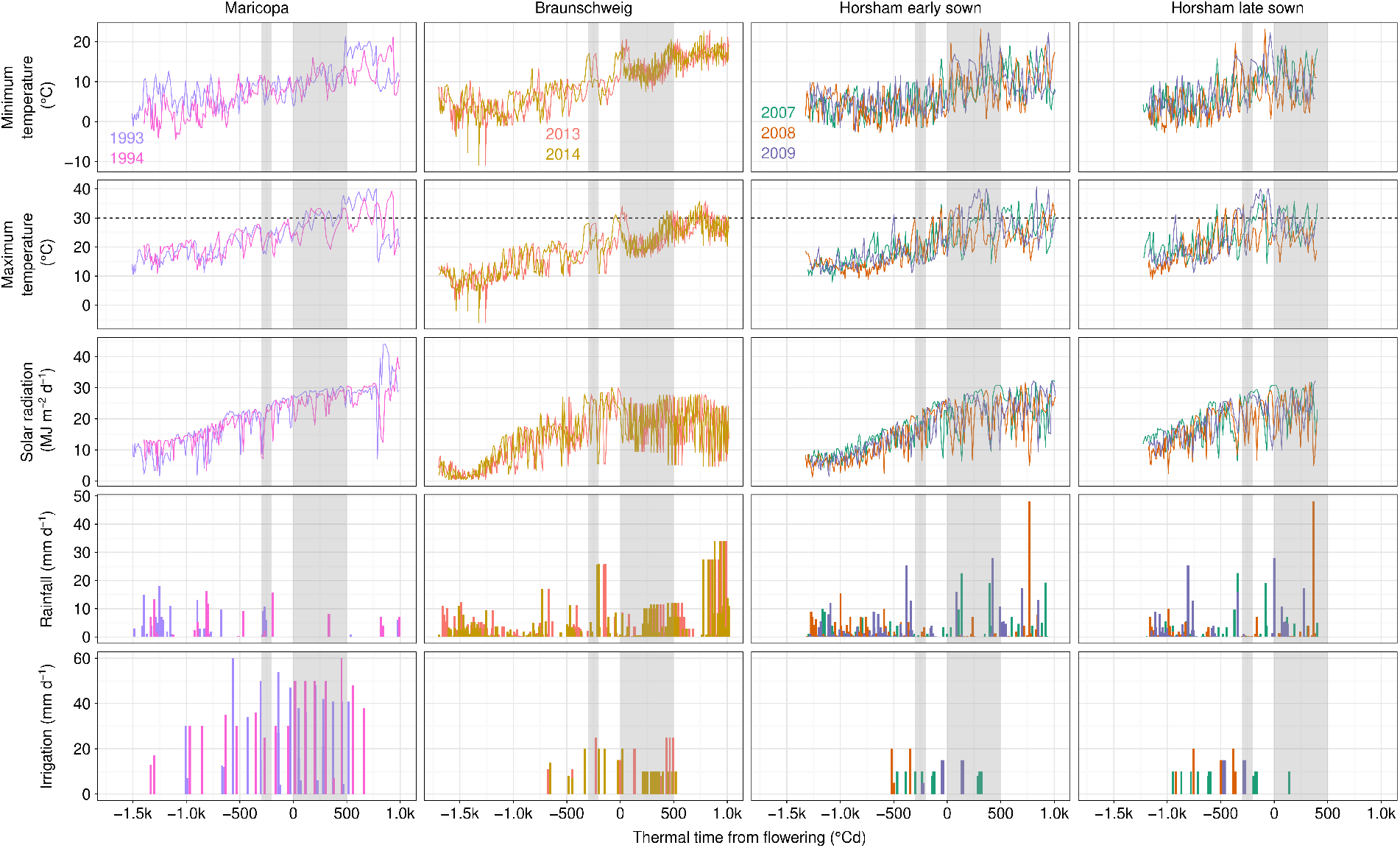
Weather conditions in Maricopa, Braunschweig, and Horsham trials as a function of thermal time from flowering. Data represent control treatments (ambient CO_2_, irrigated, early sowing) and the late sowing in Horsham. Coloured lines show minimum and maximum temperatures, and solar radiation for the studied years, while daily rainfall and applied irrigation are displayed with coloured bars. The data extend from sowing to maturity. Vertical dashed lines correspond to flowering dates for each experimental year. Grey shaded areas represent the pre-flowering (-300 to -200°Cd) and post-flowering (0 to 500°Cd) heat stress sensitivity windows defined by Ullah et al. (2024). The horizontal dashed line in the maximum temperature panels marks the 30°C threshold.

**Figure 2:**
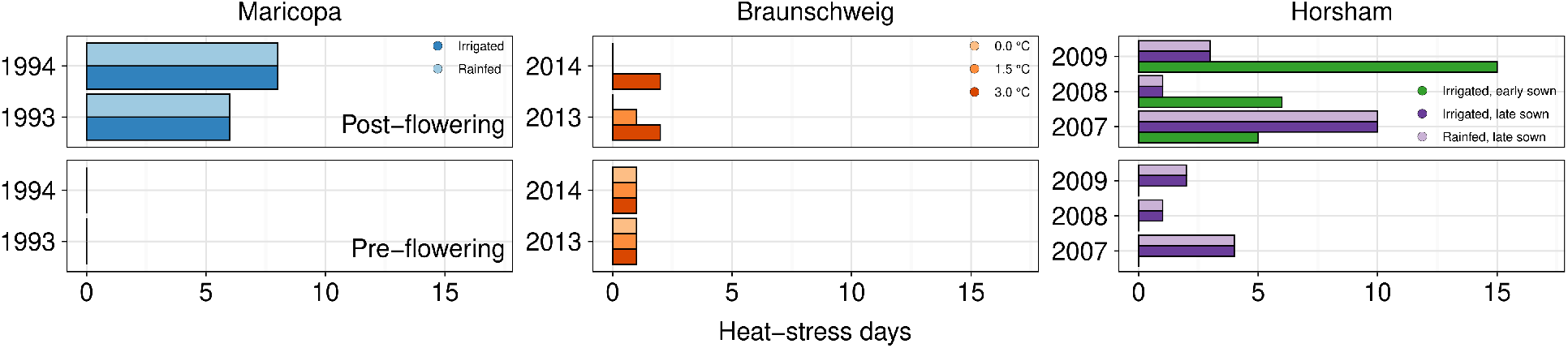
Pre-(first row) and post-flowering (second row) heat-stress exposure for each trial and treatment. Heat-stress exposure was expressed as the number of days exceeding 28°C during the pre-flowering critical stage and the number of days above 32°C post-flowering, using the thermal windows and thresholds defined by Ullah et al. (2024) and illustrated in Figure 1.

Conversely, in Maricopa, where no significant heat stress was reported, daily maximum temperatures were substantially higher and likely imposed a severe heat stress on developing grains, frequently reaching and sometimes exceeding 40°C. Specifically, trials in Maricopa experienced 16 and 10 days with temperatures above 32°C between 0 and 500°Cd after flowering during the maximum grain length phase in 1993 and 1994, respectively, across all CO_2_ and water treatments (Figs. 1 and 2).

Horsham trials also displayed considerable heat stress, particularly for the early-sown treatments (Figs. 1 and 2). In 2009, early-sown wheat at Horsham was exposed to (i) 2 days above 28°C during the critical pre-flowering phase (-300 to -200°Cd from flowering) identified by Ullah et al. (2024) as particularly sensitive for grain number determination under heat stress, and to (ii) 13 days above 32°C during the post-flowering phase and 2 days during pre-flowering (Fig. 2). Similarly, in 2007 and 2008, early-sown crops experienced respectively 5 and 6 days above this threshold during grain development. In contrast, late-sown treatments generally faced lower heat stress during grain filling, with 0 to 6 days exceeding 32°C, although they were often exposed to more frequent heat stress during the flowering phase (up to 3 days) compared to the other trials (Fig. 2). A notable exception occurred in 2007, where the early-sown crop largely escaped a period of high maximum temperatures due to its earlier developmental timing, while the late-sown crop was exposed to it during grain filling — resulting in higher heat stress for the late-sown treatment than for the early-sown, contrary to the intended experimental design. This highlights that using sowing date as a proxy for thermal treatment introduces uncontrolled variability across years, and underscores the value of multi-year experiments for robustly characterising thermal environments.

Water-stress patterns, as indicated by the simulated APSIM water stress index (WSI), revealed important insights into the actual growing conditions experienced across sites (Fig. 3). We acknowledge that using APSIM-derived indices to characterise stress conditions while simultaneously evaluating APSIM introduces a degree of circularity; however, no independent stress measurements were available across all treatments, and this approach is consistent with common practice in model evaluation studies where simulated indices serve as proxies for unobserved physiological states. At Maricopa, water stress was present even in the supposedly well-watered control (irrigated) treatment, with WSI declining to approximately 0.6 during mid-grain filling (near 500°Cd after flowering) in 1993. In contrast, the 1994 control treatment experienced minimal water stress throughout the season, highlighting substantial inter-annual variability in water availability. Rainfed treatments experienced considerably more severe water limitations, with WSI dropping to 0.3 during the critical pre-flowering period. Notably, crops grown under elevated CO_2_ (eCO_2_) consistently exhibited higher WSI values (reduced stress) compared to ambient CO_2_ at the same water treatment level, reflecting the physiological effect of reduced stomatal conductance under eCO_2_, which decreased transpiration demand and thus alleviated water stress (Table 4, Fig. 3).

**Figure 3:**
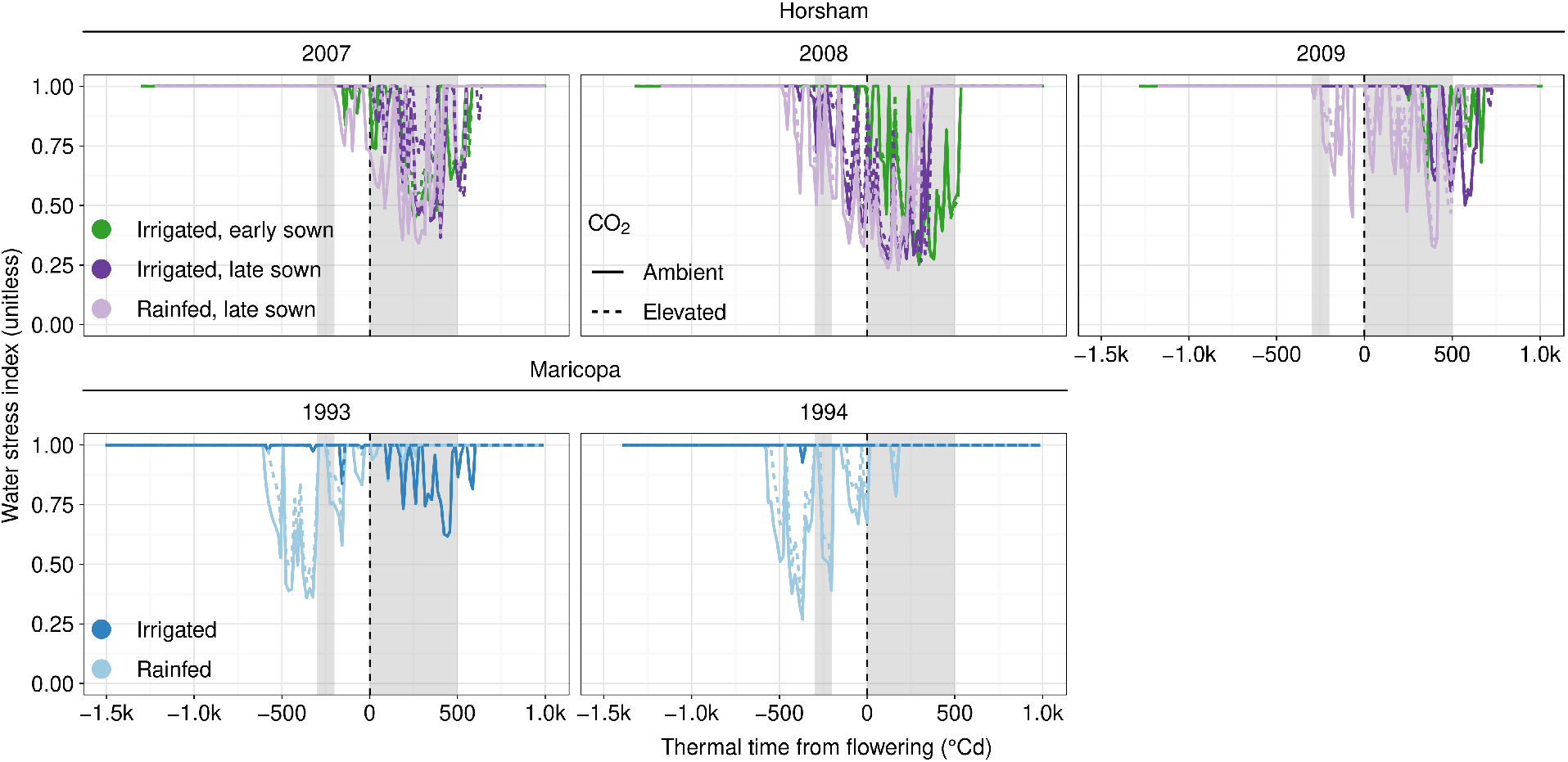
Water stress index (WSI), simulated as the supply-to-demand ratio using APSIM, for Horsham and Maricopa across all treatments. WSI values range from 1 (no water limitation) to 0 (no water available to the crop). Data for Braunschweig are not presented because WSI values remained at 1 throughout the crop growth cycle for all treatments. Grey shaded areas represent the sensitivity windows defined by Ullah et al. (2024) as shown in Figure 1.

At Horsham, simulated WSI revealed marked water stress across all treatments, including the control irrigated conditions which were designed to eliminate water limitation. For early-sown crops, water stress started before flowering and persisted through mid-grain filling across all three experimental years (2007, 2008, 2009). The severity of this stress varied inter-annually, with 2008 experiencing the most extreme water limitation (WSI declining to 0.25), indicating that even irrigated treatments were subjected to substantial drought stress (Fig. 3). This finding has potential implications for model calibration, as the control treatments used for parameterisation at Horsham were themselves affected by water stress, meaning the models were not calibrated under truly optimal, non-stressed conditions.

In contrast to Maricopa and Horsham, Braunschweig did not appear to experience any water stress throughout the growing season across all treatments. WSI remained at 1.0 for all conditions, indicating that water was never a limiting factor at this site (data not shown in Fig. 3 due to constant values).

### 3.2 Simulations of control treatments (used to parameterise the cultivars)

Both APSIM and STICS were first calibrated for the three studied cultivars using data from the control treatments at the three studied locations. For both models, simulations of LAI and above-ground biomass dynamics closely fitted observations, both during canopy growth and senescing phases (Fig. 4). Similarly, simulated data of phenology, yield and its components closely reflected observations. The final set of calibrated parameters are shown in Supplementary Figure S1.

**Figure 4:**
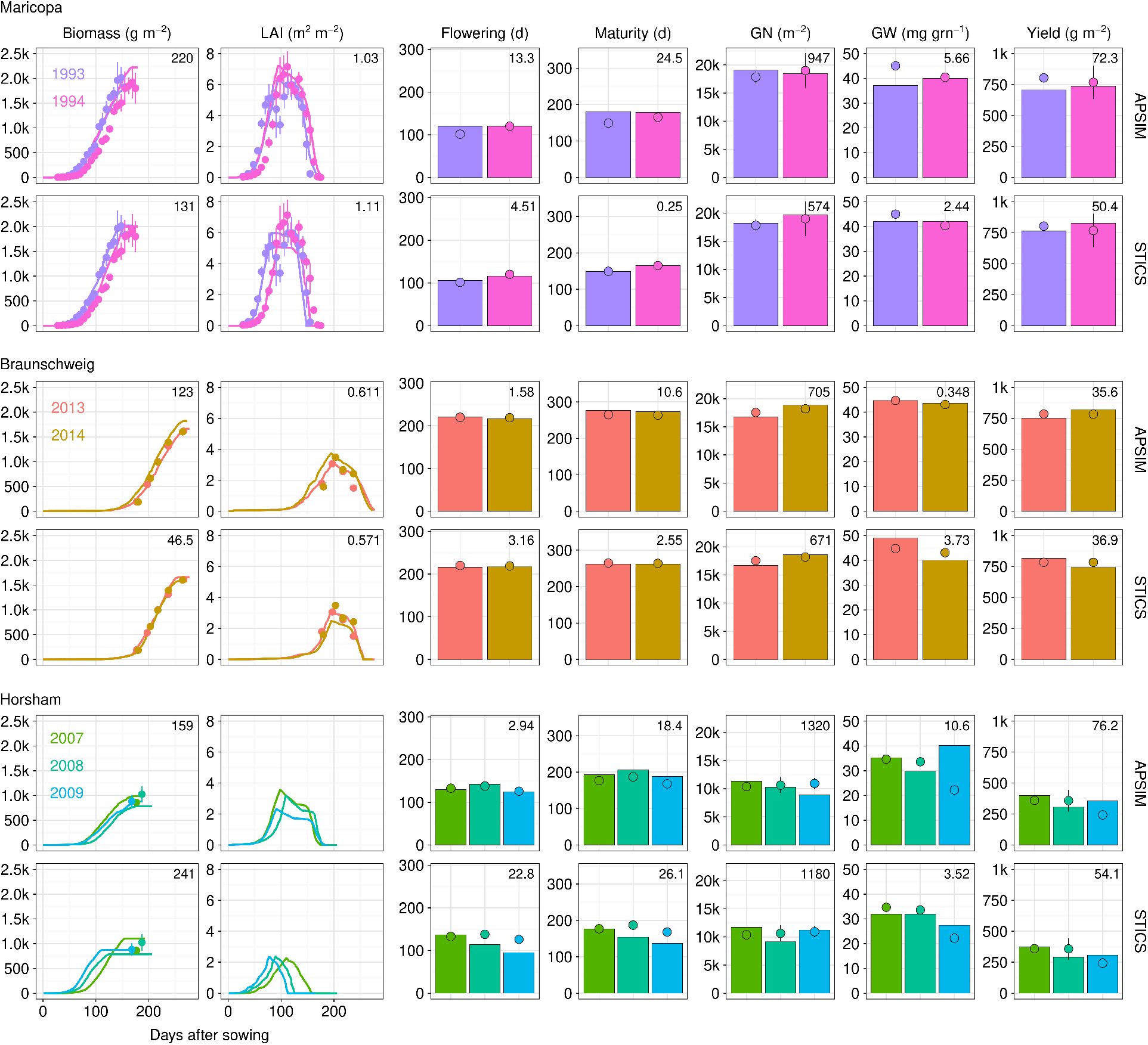
Calibration results of APSIM and STICS models against observations from control treatments of Maricopa, Braunschweig, and Horsham trials. The two left columns illustrate the dynamics of biomass and leaf area index (LAI) for the control treatment of each trial, with simulations (lines) and observed data (points). The subsequent plots to the right present flowering, harvest time (days from sowing), grain number (GN), individual grain weight (GW), and yield for simulations (bars) and observations (points). Error bars represent standard errors. No LAI measurements were conducted at Horsham. The values in the top-right corners indicate the root mean squared error (RMSE) between simulated and observed data.

### 3.3 Model evaluation under contrasting atmospheric CO_2_, soil water and air temperature conditions

Following cultivar calibration, the model predictive capabilities were tested under different conditions of atmospheric CO_2_, water stress, and thermal patterns. The simulated effects of the treatments were compared with observations across all trials and treatments (Figs. 5, 6, 7, S3, S5, and S7). Both models reproduced phenology and biomass dynamics as well as yield relatively well, with simulations generally tracking the observed effect of the treatments (RRMSE of 5–20%; Fig. 7). A notable exception was yield and biomass under rainfed conditions at Maricopa and Horsham, where STICS overestimated values relative to observations, consistent with its underestimation of water stress mentioned above. At Braunschweig, both models captured the limited yield reductions observed under moderate heat treatments (1.5°C and 3°C). At Horsham, performance was more variable across years and treatment combinations, with both models showing larger deviations from observations, particularly for grain number and yield under late-sown treatments. Overall, APSIM tended to outperform STICS in reproducing both the direction and approximate magnitude of treatment effects across sites (Fig. 6). At Horsham, the direction and magnitude of eCO_2_ effects were well captured by both models under irrigated conditions, but not under rainfed conditions. Under aCO_2_, rainfed effects were relatively well predicted in APSIM; however, neither model simulated properly the rainfed treatment under eCO_2_. For instance, APSIM captured the negative effect of the imposed drought stress but underestimated its magnitude, suggesting that interactions between eCO_2_ and water deficit are not fully resolved in this model.

**Figure 5:**
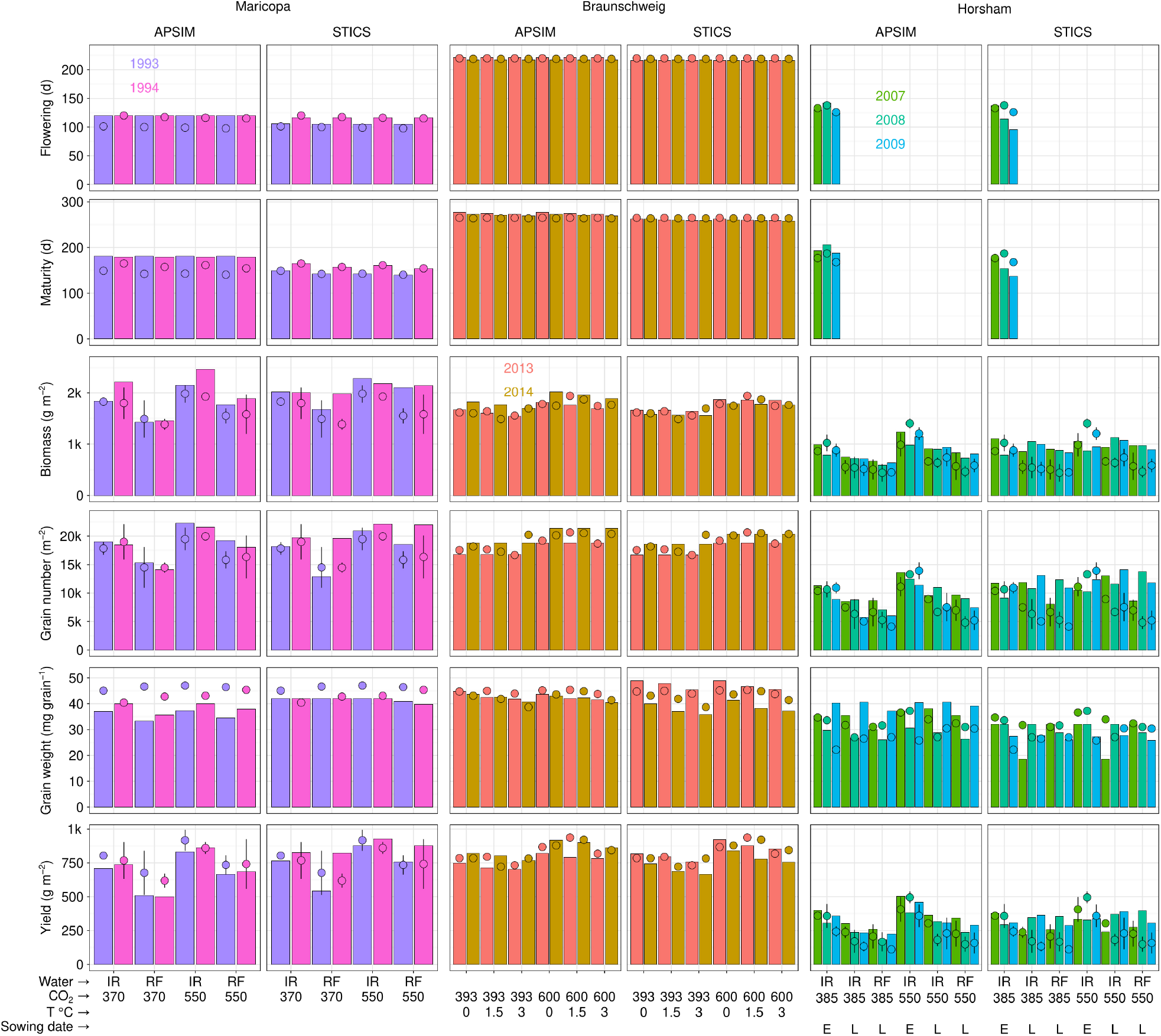
Simulated (bars) and observed (points: means +/- standard errors) treatment effects for Maricopa, Braunschweig, and Horsham trials. In Maricopa, crops were grown with two CO_2_ levels (370 and 550ppm) under two water availability conditions (IR: irrigated and RF: rainfed). In Braunschweig, similar CO_2_ levels were imposed in combination with three temperature treatments during the grain-filling period, with temperature increases of 0, 1.5, and 3°C from early to late grain filling (i.e. 13 and 5 days after flowering to 42 and 37 days after flowering, in 2014 and 2015, respectively). In Horsham, plots were subjected to similar ambient and elevated CO_2_ levels while experiencing different water and temperature patterns, with warmer temperatures occurring for the late-sown crops (sowing date E: early, L: late). In Horsham, phenology data was available for only the control treatments evaluated. Relative effects compared to the control treatments are presented in Figure 6.

**Figure 6:**
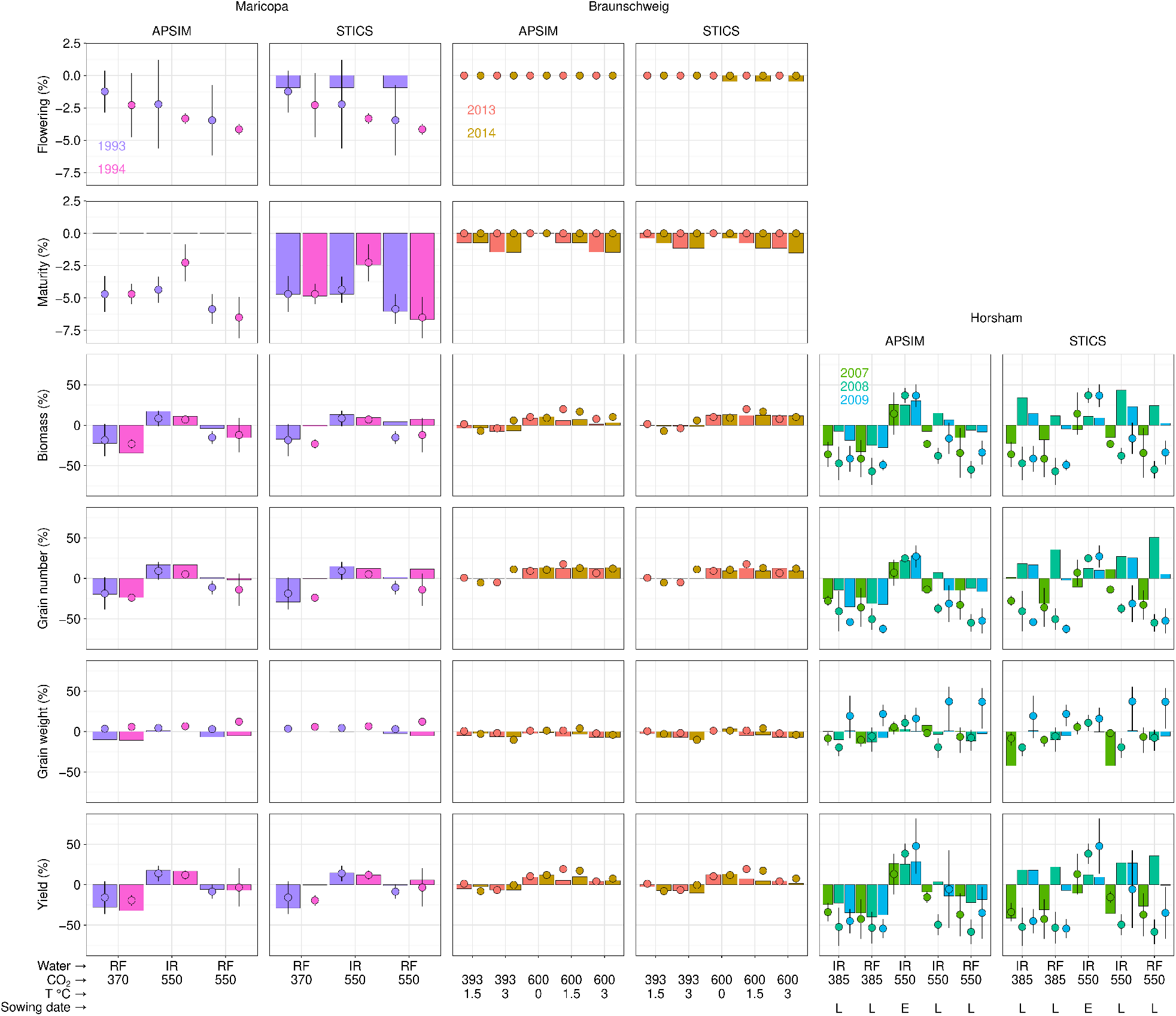
Relative simulated (bars) and observed (points) effects of all studied treatments compared to their respective control treatments in all trials. Relative effects were determined as a percentage, i.e. as (value - control) / control * 100. Since only control data was available for phenology in Horsham, relative effects for flowering and maturity could not be computed in Horsham trials. Effects (in absolute value) are presented in Figure 5.

**Figure 7:**
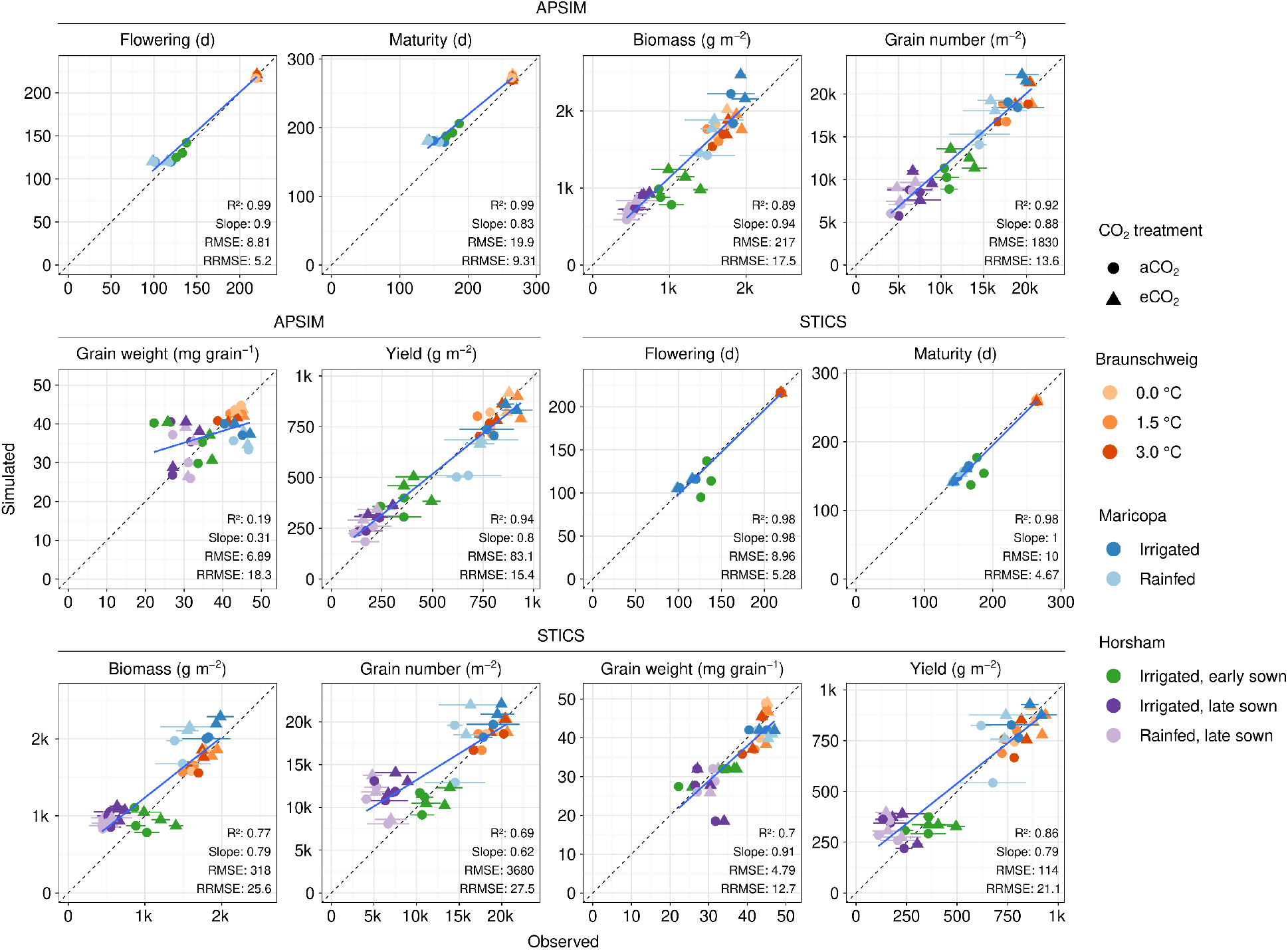
Simulated against observed days to flowering and maturity, biomass, grain number, individual grain weight and yield in all trials and treatments. Results from all three sites are displayed together, with different colour scales representing different treatments. For each trait and each model, a solid blue line represents the linear regression between simulated and observed data, while the dashed line corresponds to the 1:1 line. Regression statistics, including R^2^, slope, as well as root mean squared error (RMSE) and relative RMSE (RRMSE, % of mean observed value) are presented in the bottom-right corner of each graph. Note that RMSE and RRMSE quantify the overall difference between simulated and observed values, independent of the regression analysis.

Overall, both models achieved high accuracy for phenology (APSIM: R^2^=0.99, RRMSE=5.2% for flowering; STICS:R^2^=0.98, RRMSE=5.3%; Fig. 7). For yield, APSIM (R^2^=0.94, RRMSE=15.4%) outperformed STICS (R^2^=0.86, RRMSE=21.1%).The most striking divergence between models was for grain number and grain weight: APSIM captured grain number well (RRMSE=13.6%) but tended to simulate grains with constant weight in many environments, resulting in low prediction accuracy (R^2^=0.19, slope=0.31), whereas STICS showed the opposite pattern with poor grain number prediction (R^2^=0.69, RRMSE=27.5%) but reasonable individual grain weight estimations (R^2^=0.70, RRMSE=12.7%).

### 3.4 Model performance in different types of environmental conditions

To evaluate the model performance in relation to specific environmental pressures, simulations were categorised into six stress configurations based on heat and drought stress(es) that crops experienced pre-flowering and post-flowering. Crops were categorised as (i) heat stressed (“Heat”) if exposed for at least 2 days at >28°C during the pre-flowering critical phase or at least 2 days at >32°C post-flowering, as (ii) water stressed (“Water”) if experiencing a WSI < 0.7 pre-flowering (-300 to 0°Cd) or post-flowering (0 to 500°Cd), and (iii) both heat and water stressed (“Both”) when both stresses occurred during the respective pre- or post-flowering periods. The “pre- | post-flowering” categories that occurred in the tested trials were: “None | None”, “None | Heat”, “None | Both”, “Heat | Both”, “Water | Heat”, and “Both | Both” (Figs. 8 and S8). The dataset was unevenly distributed across categories, with “None | None” being the most represented (n = 12), followed by “None | Both” (n = 9) and “Both | Both” (n = 6), while “None | Heat”, “Water | Heat”, and “Heat | Both” were less represented (n = 3–4 each), reflecting the relative scarcity of single-stress conditions in the available FACE trials. In addition, in all categories, only trials from a single site were represented, which is quite restrictive (Fig. S8). Relative root mean square error (RRMSE) was calculated for all variables within each category and CO_2_ levels to assess the model prediction accuracy in those situations (Fig. 8).

**Figure 8:**
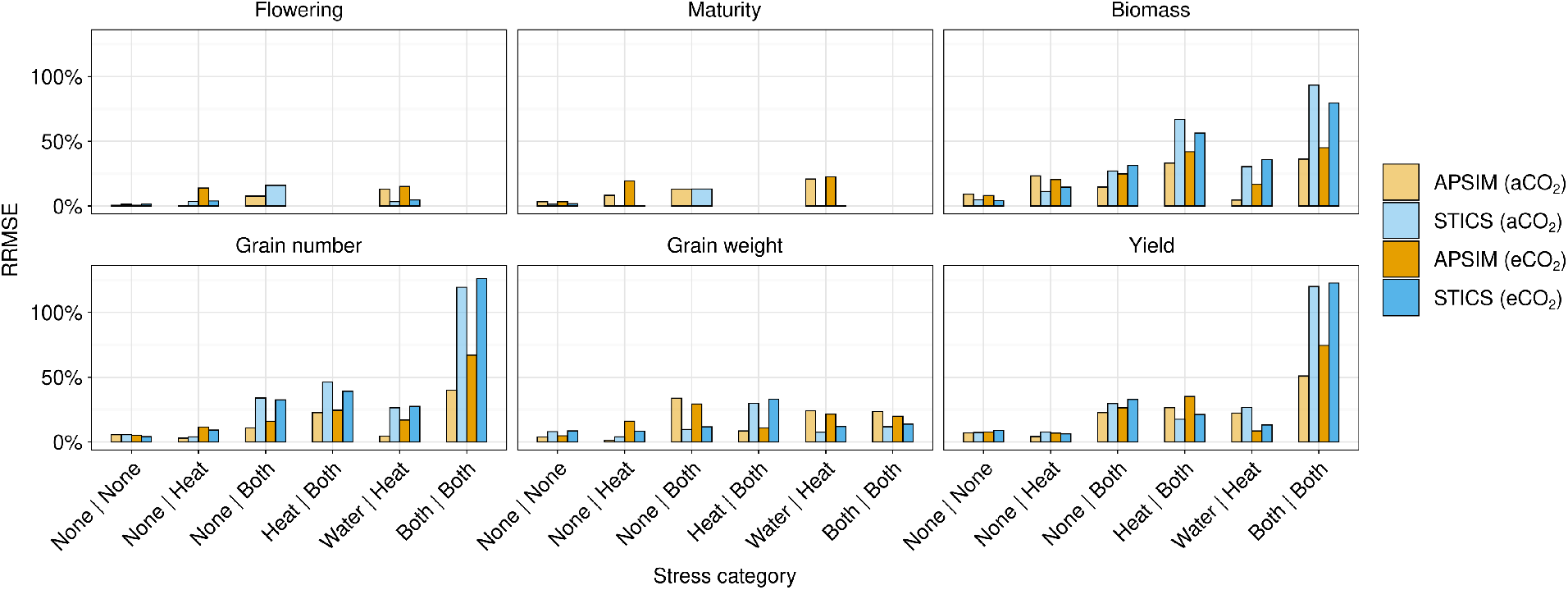
Relative root mean square errors (RRMSE) calculated by stress category, model and CO_2_ level. Stress categories follow Ullah et al. (2024), defined by simulated heat exposure (days >28°C between -300 to -200°Cd for heat or -300 to 0°Cd for water before flowering or >32°C between 0 and 500°Cd after flowering) and water stress (WSI < 0.7) within these respective periods, irrespective of the atmospheric CO_2_ level. Categories [Pre-flowering | Post-flowering] include: “None | None” (no stress); “None | Heat” (post-flowering heat); “None | Both” (post-flowering combined); “Heat | Both” (pre-flowering heat, post-flowering combined); “Water | Heat” (pre-flowering water, post-flowering heat); and “Both | Both” (combined stress in both periods). Only categories present in the dataset are displayed. Notably, categories with repeated stress of the same type across both phases (e.g., Water | Water or Heat | Heat) were absent from the dataset, which prevents a clean separation of stress temporality from stress combination in the analysis.

The lowest errors were associated with conditions of minimal stress, typical of treatments where water availability was non-limiting and heat exposure was low (Fig. S8). Under these low-stress conditions (None | None), RRMSE for yield ranged from 6.9% to 7.1% under ambient CO_2_ for both models, but slightly increased to 7.7–8.9% under elevated CO_2_, demonstrating that while APSIM and STICS achieve high accuracy when environmental drivers are favourable in both aCO_2_ and eCO_2_ (Figs. 8 and S8).

Under stress, prediction errors generally increased according to the type of stresses (Both > Heat > Water) and their timing (pre-flowering > post-flowering), often exacerbated by elevated CO_2_ when there is a water stress. Across situations, errors in predicting grain number due to a pre-flowering stress had massive repercussions on the accuracy of yield predictions. By contrast, the prediction accuracy of individual grain weight remained relatively modest in comparison, despite being influenced by both the number of grains set and the occurrence of post-flowering stress affecting grain filling.

The impact of pre-flowering water stress on grain number (the main driver of wheat yield) was relatively well simulated across all simulated crops. However, prediction errors increased with stress complexity, being lowest under water stress alone, intermediate under heat stress, and highest under combined water and heat stress (Water < Heat < Both), with RRMSE values of 4.3, 22.6, and 39.9%, respectively, for APSIM under aCO_2_ (Fig. 8). Under eCO_2_, prediction errors for grain number increased substantially when crops experienced water stress particularly in APSIM, with RRMSE rising from 16.8 to 24.5 for Water | Heat and to 67% for Both | Both. These errors strongly affected yield prediction accuracy. Notably, in the studied trials, no crops experienced pre-flowering stress without also being exposed to post-flowering stress.

The impact of post-flowering stress alone (i.e. not preceded by pre-flowering stress) on yield could not be evaluated for water stress due to the absence of such cases. Errors remained relatively low under post-flowering heat stress alone (None | Heat; RRMSE of 4.0–7.8%) but increased markedly under combined stresses (None | Both; RRMSE of 22.6–29.6%) under aCO_2_. Elevated CO_2_ had no substantial effect on RRMSE for post-flowering heat stress alone (None | Heat), but it increased prediction errors when both stresses occurred (None | Both) in both models, from 22.6–29.6% under aCO_2_ to 26.3–32.7% under eCO_2_. A similar pattern was observed when pre-flowering heat stress preceded combined post-flowering stress (Heat | Both), with RRMSE increasing from 17.5–26.4% under aCO_2_ to 21.2–35.0% under eCO_2_.

The most extreme prediction errors were consistently associated with simultaneous water and heat stress occurring both before and after flowering (Both | Both; Figs. 8 and S8). Under these multi-stress conditions, eCO_2_ further degraded model performance, particularly in APSIM, where RRMSE increased from 50.9% to 74.5%. For STICS, errors were and remained extreme above 120% regardless of CO_2_ level. This deterioration in prediction accuracy highlights a fundamental limitation of both models to precisely predict yield formation under complex interacting environmental stresses, at least for the trials we tested here (i.e. late-sown Horsham trials).

## 4 Discussion

### 4.1 Confounding effects of unintended stress on calibration and validation

A critical finding of this study is the substantial divergence between intended experimental treatments and the actual environmental conditions experienced by the crops, underscoring the need to carefully characterise environments to which crops are exposed (Chenu, 2015). The presence of significant heat and water stress in treatments intended as non-limiting controls at Maricopa and Horsham (Figs. 1 and 3) challenged the traditional approach to crop model evaluation, which typically includes non-limiting conditions to determine genotype-specific parameters. Our analysis revealed that control treatments at these trials were themselves affected by stress, meaning that, in some cases, cultivar parameters were fitted to crops already undergoing physiological adjustments to environmental limitation. This challenges the proper calibration of traits such as radiation-use efficiency (RUE), with errors that may propagate through the model via compensatory adjustments of other parameters, potentially introducing errors in the simulation of associated processes (Gawinowski et al., 2025a). Such confounding effects, possibly exacerbated by the relatively limited number of observations available, may explain why both APSIM and STICS exhibited higher baseline RRMSE of cv. Yitpi at Horsham (17–24%) compared to cv. Batis which experienced truly stress-free conditions at Braunschweig (RRMSE of 3–5%). This also suggests that part of the apparent success of the models in simulating stress treatments may arise from calibration under already similar stressed conditions, rather than accurately simulating the underlying physiological response to a deviation from potential growth.

### 4.2 Challenges in simulating water, heat and combined stress

The hierarchical escalation of prediction errors from single stresses to combined water and heat stress suggest the presence of a limitation in current process-based models. Both APSIM and STICS generally handle individual stressors tested here with reasonable accuracy, except for water stress in Maricopa with STICS, which effect was strongly underestimated likely due to the underestimation of evapotranspiration. Similar issues were actually reported in other modelling studies for this specific trial with CERES-Wheat (Tubiello et al., 1999), LINTULCC2, AFRWHEAT2 and Sirius (Ewert et al., 2002). However, in the present study, APSIM managed to simulate water stress effects accurately in Maricopa, suggesting the relevance of the Penman-Monteith method to compute evapotranspiration with the energy balance, rather than the Shuttleworth and Wallace approach in this case. This comparison raises interesting perspectives to improve the simulation of evapotranspiration in some crop models. Webber et al. (2025) pointed out that drought stress is indeed often underestimated in crop models in semi-arid conditions, and they revealed a systematic bias with an underestimation of evapotranspiration, even for the median of the model ensemble.

The impact of pre-flowering heat stress on grain number could also be better represented in both APSIM and STICS (Fig. 8). Notably, these models currently do not simulate the direct effects of heat stress on processes affecting grain set, such as pollen meiosis. Several approaches have been proposed to represent such mechanisms and other heat-related responses in crop models (e.g. Ababaei and Chenu, 2020; Bell et al., 2016; Richetti et al., 2025), but they have yet to be rigorously evaluated. In the present study, the T-FACE treatments implemented at Braunschweig to increase post-flowering temperature using ceramic heaters did not result in any realised heat stress. Post-flowering heat stress in the absence of pre-flowering stress was therefore evaluated only for one year at Maricopa under ambient CO_2_ and for two years under elevated CO_2_ (Fig. S8). Evaluating the models using additional datasets that encompass a broader range of heat wave conditions would be valuable to more adequately assess their capacity to simulate post-flowering heat stress responses. This limitation is of particular concern given the increasing importance of heat stress under climate change.

STICS and APSIM performance deteriorated sharply when both drought and heat stress co-occurred (Fig. 8). This failure is likely rooted in the additive or multiplicative nature of current stress functions, which may not capture the synergistic physiological interactions that emerge under multi-stress occurrences. As reviewed by Richetti et al. (2025), these interactions often result in yield losses that far exceed the sum of individual stress factors, challenging the assumptions central to many current model architectures. Overall, little is known about the quantitative implications of acclimation, or about how stress effects are modified when stresses persist over different durations or co-occur with other stresses. In addition, key processes remain inadequately modelled. For instance, water stress induces stomatal closure, reducing transpirational cooling and potentially raising canopy temperatures significantly above ambient air temperature (Wu et al., 2016). The discrepancy between meteorological climate and the actual micro-climate experienced by plant organs remains a primary source of error in process-based simulations (Richetti et al., 2025). This effect is further compounded by elevated CO_2_, which also reduces stomatal conductance and thus evaporative cooling. If so, crop models that rely primarily on air temperature as the driver for development and stress thresholds could underestimate the actual thermal stress experienced by plants under these conditions. This hypothesised “double constraint”—where drought and elevated CO_2_ potentially exacerbate heat stress—could contribute to the non-linear yield declines observed in our data that were poorly captured by the models, although we do not have direct canopy temperature measurements to confirm this mechanism. Modelling canopy temperature based on energy balance rather than reliance on ambient air temperature could improve simulation of CO_2_ × water × heat interaction (Wu et al., 2016).

### 4.3 Limitations and future research directions

The scope of this evaluation was constrained by the limited availability of high-quality multi-factorial FACE datasets, which resulted in a relatively small number of trials tested. In addition, model evaluation required site-specific calibration at each location, as different cultivars were used across the experiments. While water stress can be considered a well-established constraint that has been widely evaluated in previous studies in models like APSIM, substantially more data are required to robustly assess model performance under heat stress, and even more so under combined heat and drought stress, especially across contrasting CO_2_ conditions.

The model evaluation could also be broadened in terms of CO_2_ treatments tested. The moderate CO_2_ levels (550–600 ppm) used in studied FACE trials only represent mid-century scenarios, whereas crop models are increasingly used to project crop performance over longer time horizons, including toward the end of the century, when atmospheric CO_2_ concentrations may exceed 800 ppm (Allen et al., 2020). In addition, results observed in open-air FACE systems may not fully reflect future crop responses, as fluctuations in wind speed and direction can generate rapid short-term variability in CO_2_ concentration, which contrasts with the stable atmospheric CO_2_ concentration experienced by crops in natural conditions (Allen et al., 2020).

Furthermore, models like STICS and APSIM typically operate at a daily time step, meaning that effects of rapid environmental fluctuations are not directly captured by these models. Moving beyond the typical daily resolution to incorporate sub-daily thermal dynamics could improve simulations by more accurately accounting for the duration and intensity of short-lived extreme events, such as heat waves (Richetti et al., 2025; Ullah et al., 2024).

Overall, there is a clear need for detailed controlled-environment and field studies that investigate the quantitative impact of varying stress intensities, timings/durations, and stress combinations on elementary physiological processes (e.g. organ expansion rates, stomatal conductance, senescence). Such insights and datasets would enable more rigorous testing of model algorithms and the development of improved stress responses.

## 5 Conclusion

This work highlights the critical importance of comprehensive environmental characterisation when interpreting experimental studies. Detailed analysis revealed that actual environmental conditions frequently diverged from intended experimental treatments: heat stress was more severe at sites where it was not intended (Maricopa and Horsham) than at the site where it was applied as a targeted treatment (Braunschweig), and water stress occurred even in irrigated control conditions at Maricopa and Horsham. These findings demonstrate that the distinction between “control” and “stressed” treatments can be less clear-cut than experimental designs imply. Consequently, the “control” treatments used in this study for the calibration of genotype-specific parameters were not truly stress-free as initially thought.

The analysis of prediction errors across stress categories revealed that models simulated single stressors with relatively reasonable accuracy within the scope of this evaluation, although improvement are required for water stress simulations in STICS, and for heat stress in both models. More critically, the performance of the models deteriorated sharply under combined heat and water stress. This escalation in prediction error suggests that current stress functions are insufficient to capture key synergistic physiological interactions, such as reduced transpirational cooling under drought with or without elevated CO2 exacerbates heat stress through reduced stomatal conductance. To improve the predictive capacity of crop models, future research should move towards more mechanistic modelling of stress responses and their interactions, supported by experimental designs that provide clear, quantitative response norms to environmental factors. Only through such mechanistic development can crop models deliver robust predictions in novel environments and for the correct physiological reasons, particularly in the context of increasingly complex stress conditions associated with climate change.

## Acknowledgements

The authors thank Bangyou Zheng for their valuable assistance in compiling the model process descriptions presented in Table 4. During the preparation of this manuscript, large language model (LLM) tools were used to assist with revising the text. The authors reviewed and edited all AI-assisted content and take full responsibility for the accuracy and integrity of the published work.

The authors also acknowledge support from the project REGARD (“RechErche d’analoGues climAtiques pour sélectionneR Demain”) funded by the “Fonds de Soutien à l’Obtention Végétale” (FSOV), and from the Australian Research Council (ARC Linkage Project LP210200723).

## Supplementary

### Supplementary Note S1: Temperature response functions in APSIM and STICS

The following equations describe the shape of the temperature response functions used in APSIM and STICS for the three processes affected by temperature in Table 4.

#### Development (thermal time accumulation)

APSIM uses a piecewise linear function:

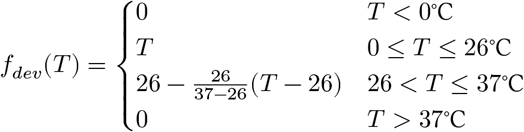

STICS uses a cardinal temperature approach with a linear increase between the base temperature (*tdmin*) and the optimum (*tdmax*), and no accumulation above the ceiling (*tcxstop*):

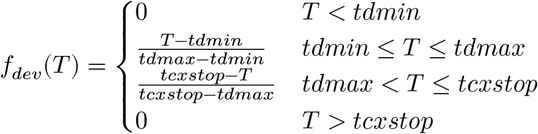

#### LAI growth

APSIM applies a temperature factor that slows growth below 12°C and ceases it at 0°C:

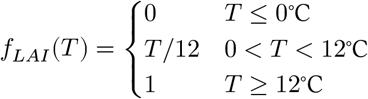

STICS uses the same cardinal temperature structure as for development, with its own set of cardinal temperatures (*temin, tcmax, tcxstop*):

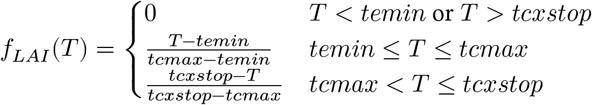

#### Radiation-use efficiency (RUE)

APSIM applies a piecewise linear multiplier:

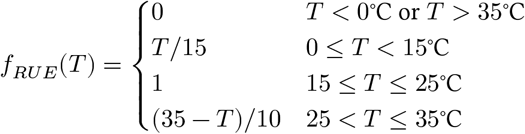

STICS uses a beta-shaped function with a plateau between the lower (*teopt*) and upper (*teoptbis*) optima:

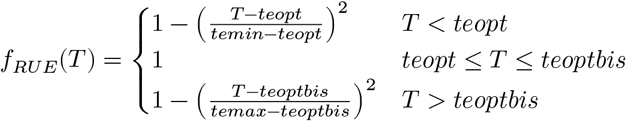

#### APSIM water deficit effect on leaf senescence

APSIM accelerates leaf senescence under water stress via a linear function of the water supply-to-demand ratio *F*_*w*_ (0= full stress, 1 = no stress):

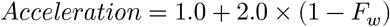

At no stress (*F*_*w*_ = 1.0), the acceleration factor is 1.0 (normal ageing rate); at maximum stress (*F*_*w*_ = 0.0), the factor reaches 3.0, meaning leaves senesce three times faster than under unstressed conditions.

## Supplementary Tables and Figures

**Supplementary Table S1:**
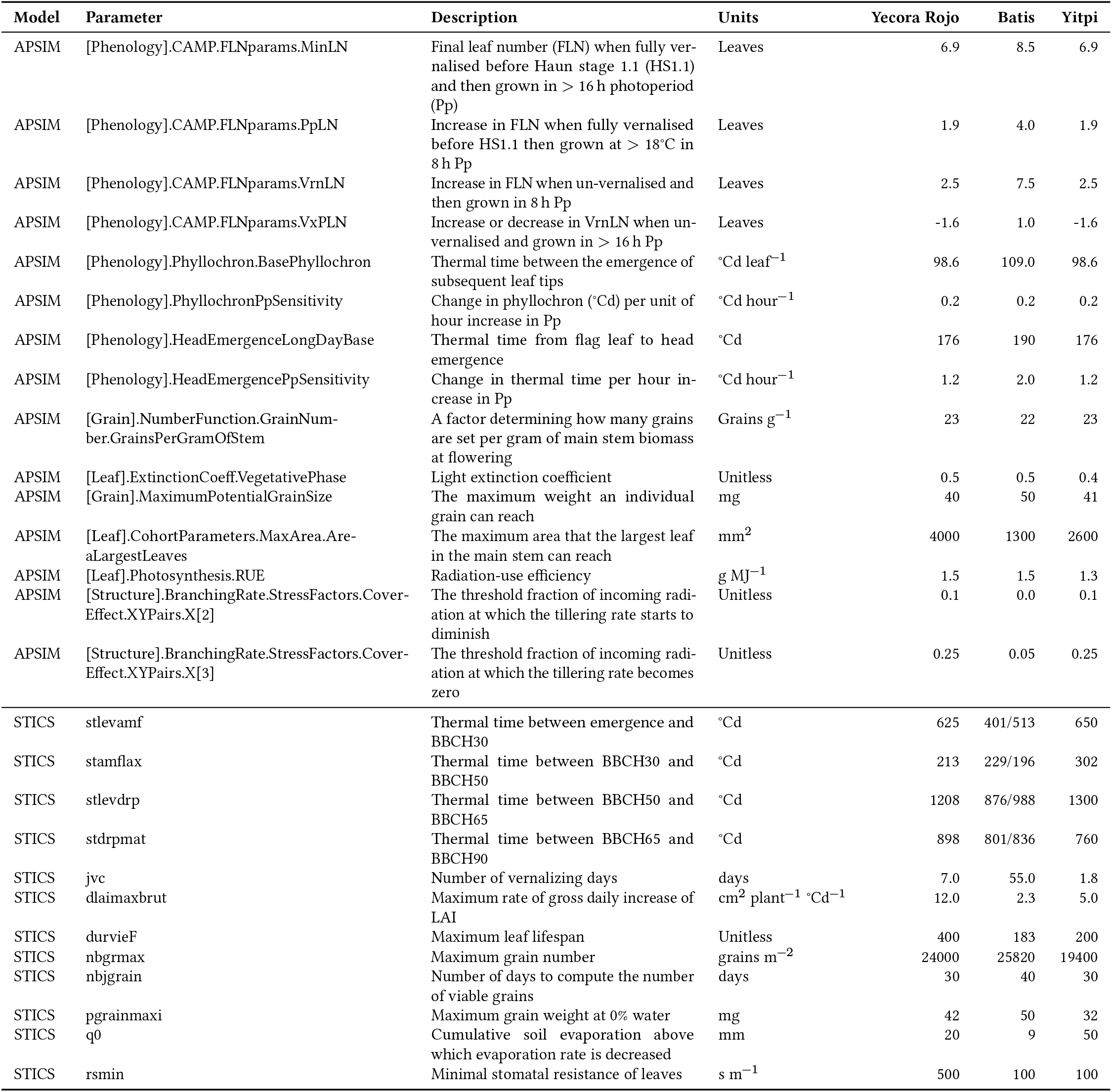
Calibrated parameter values, including APSIM and STICS names, definitions, and units. Values reflect calibrations for Maricopa (Yecora Rojo), Braunschweig (Batis), and Horsham (Yitpi). Note that STICS parameters **stlevamf, stamflax, stlevdrp**, and **stdrpmat** were calibrated independently for cultivar Batis across the 2013 and 2014 Braunschweig seasons; these values are displayed separated by a forward slash.

**Figure S1:**
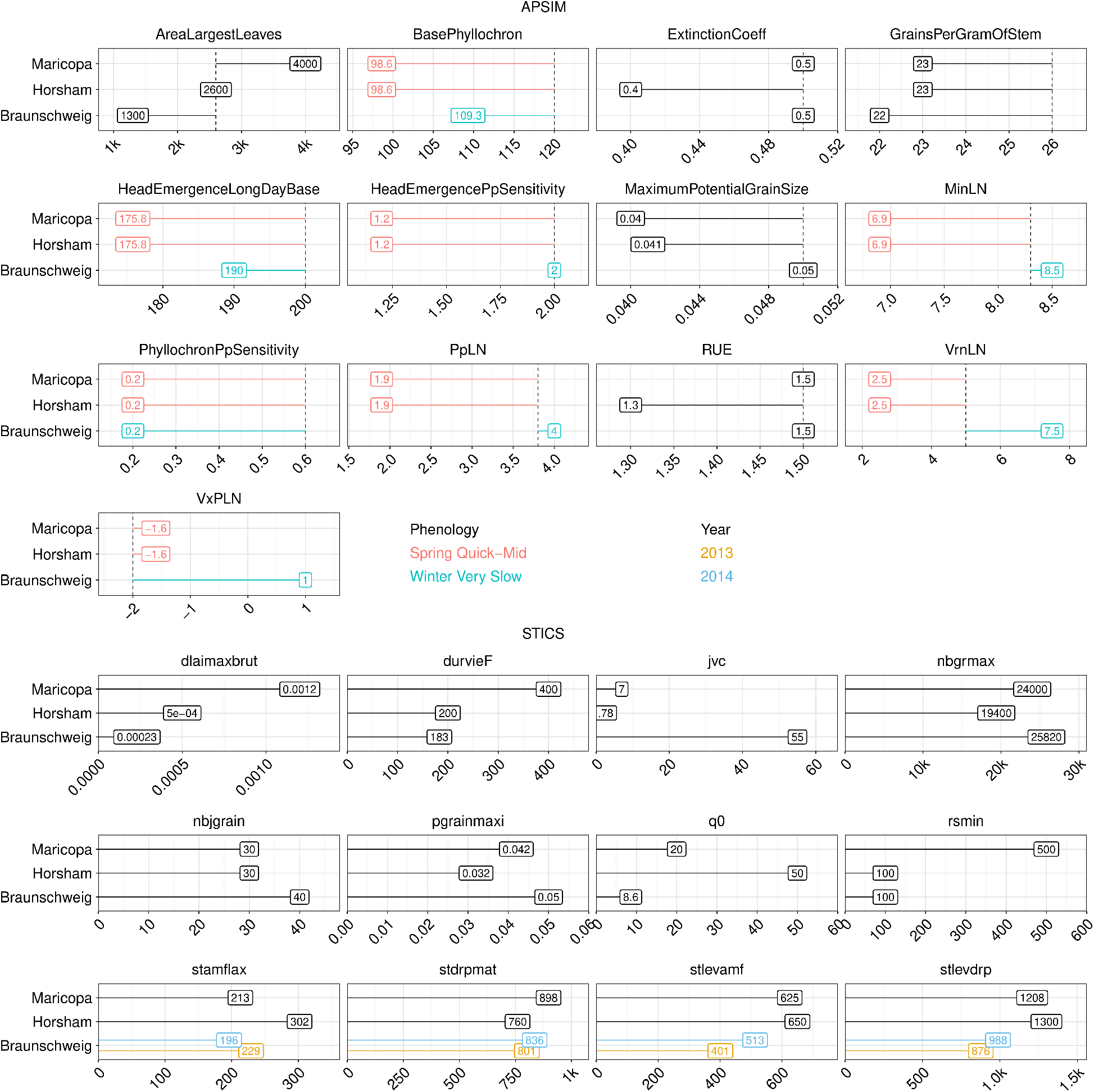
Model parameters for APSIM and STICS calibrated using control data. For APSIM, wheat phenology was established using generic cultivars from the standard distribution: Braunschweig was modelled using ‘Winter Very Slow,’ while Maricopa and Horsham used ‘Spring Quick-Mid.’ Additional parameters were adjusted site-specifically to maximise data alignment, with default APSIM values indicated by vertical dashed lines. For STICS, the parameters stamflax, stdrpmat, stlevamf, and stlevdrp were calibrated independently for the 2013 and 2014 seasons in Braunschweig. Detailed definitions for all parameters are provided in Supplementary Table S1.

**Figure S2:**
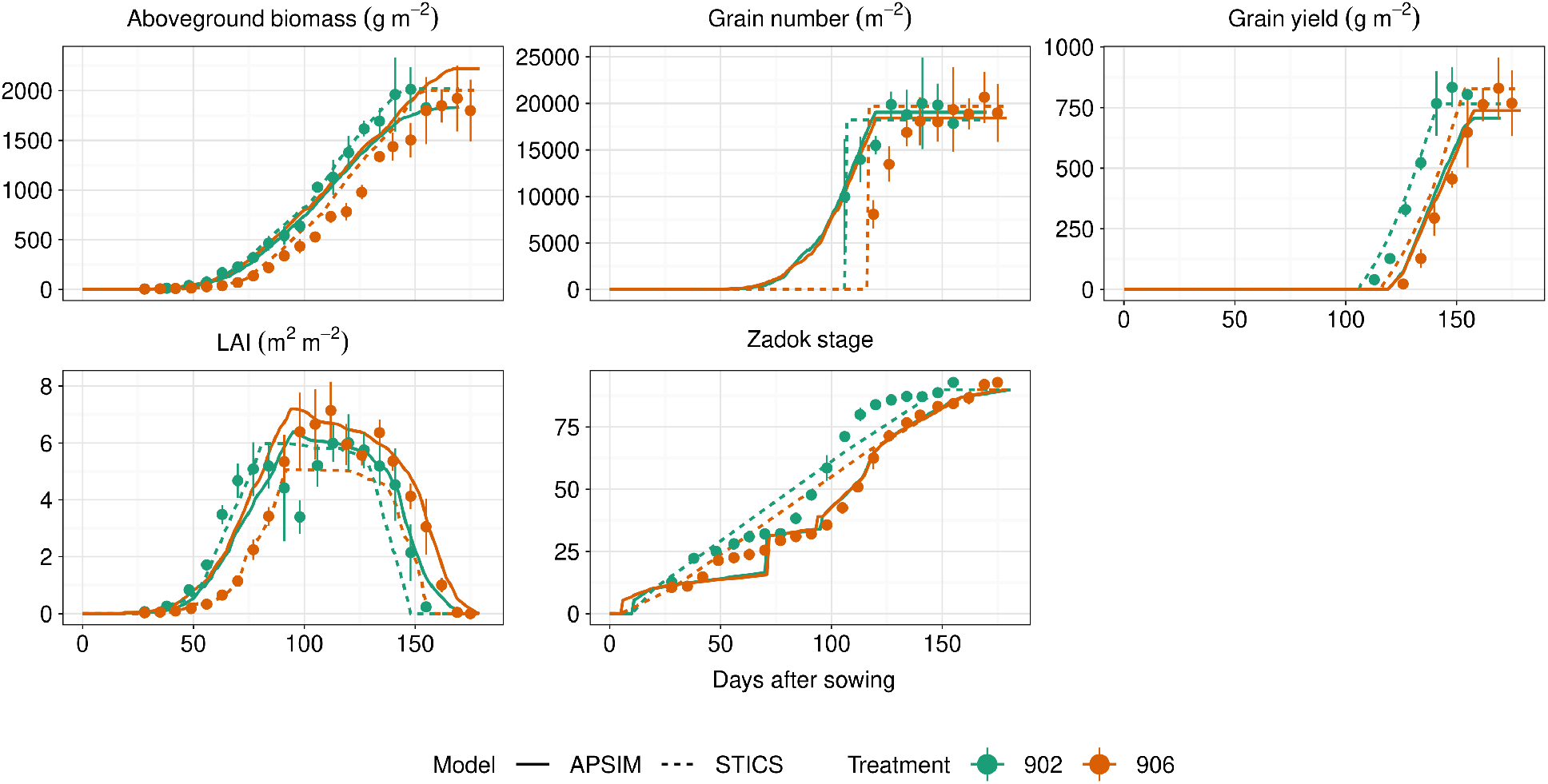
Calibration at Maricopa site using control treatments only, i.e. ambient CO_2_, no water stress. Points indicate observation means, and error bars are standard deviations.

**Figure S3:**
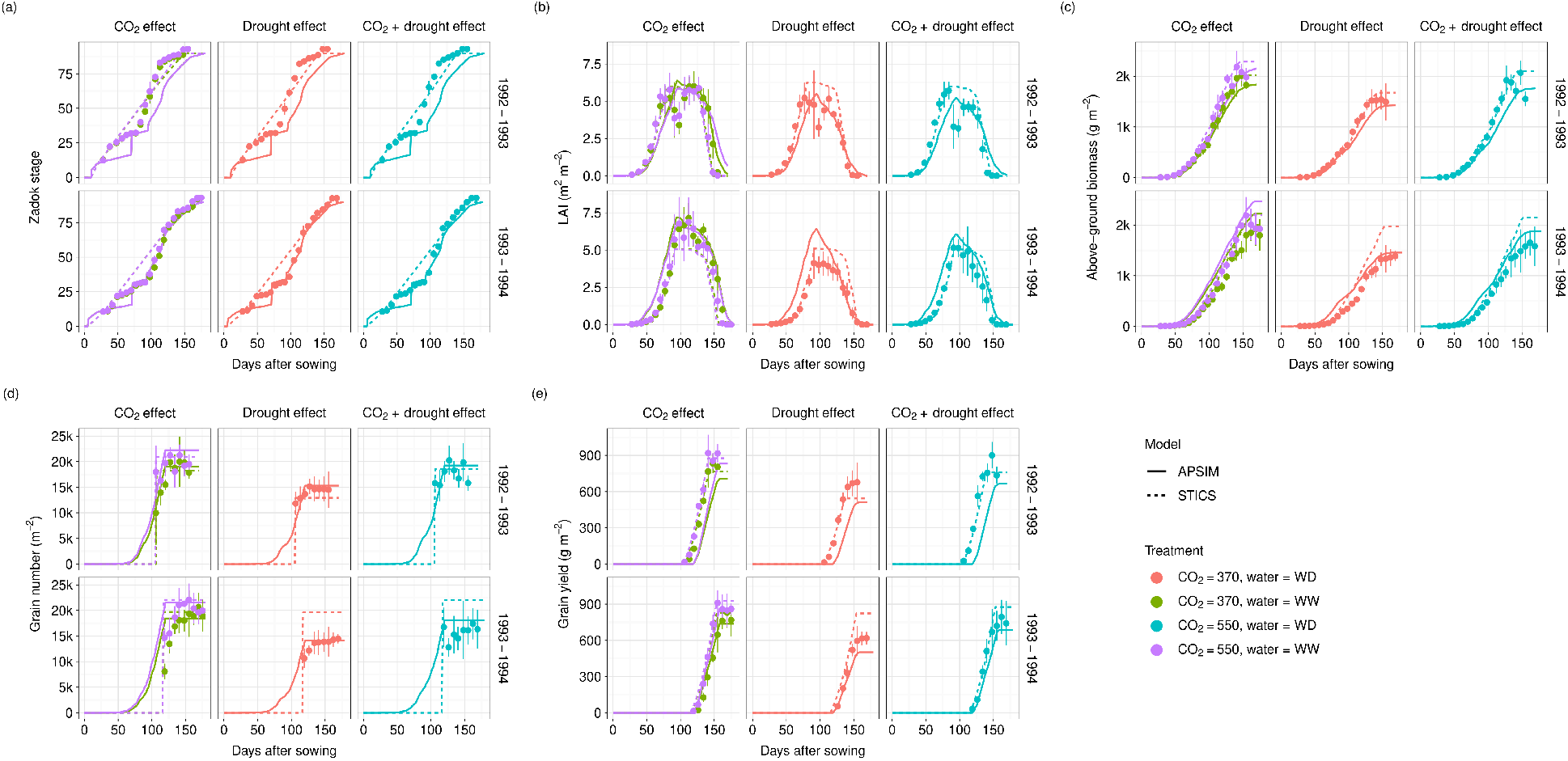
Simulated vs. observed (a) phenology (Zadok scale), (b) leaf area index (LAI), (c) above-ground biomass, (d) grain number, and (e) grain yield for all treatments in Maricopa. Points represent observed data; error bars indicate standard deviations.

**Figure S4:**
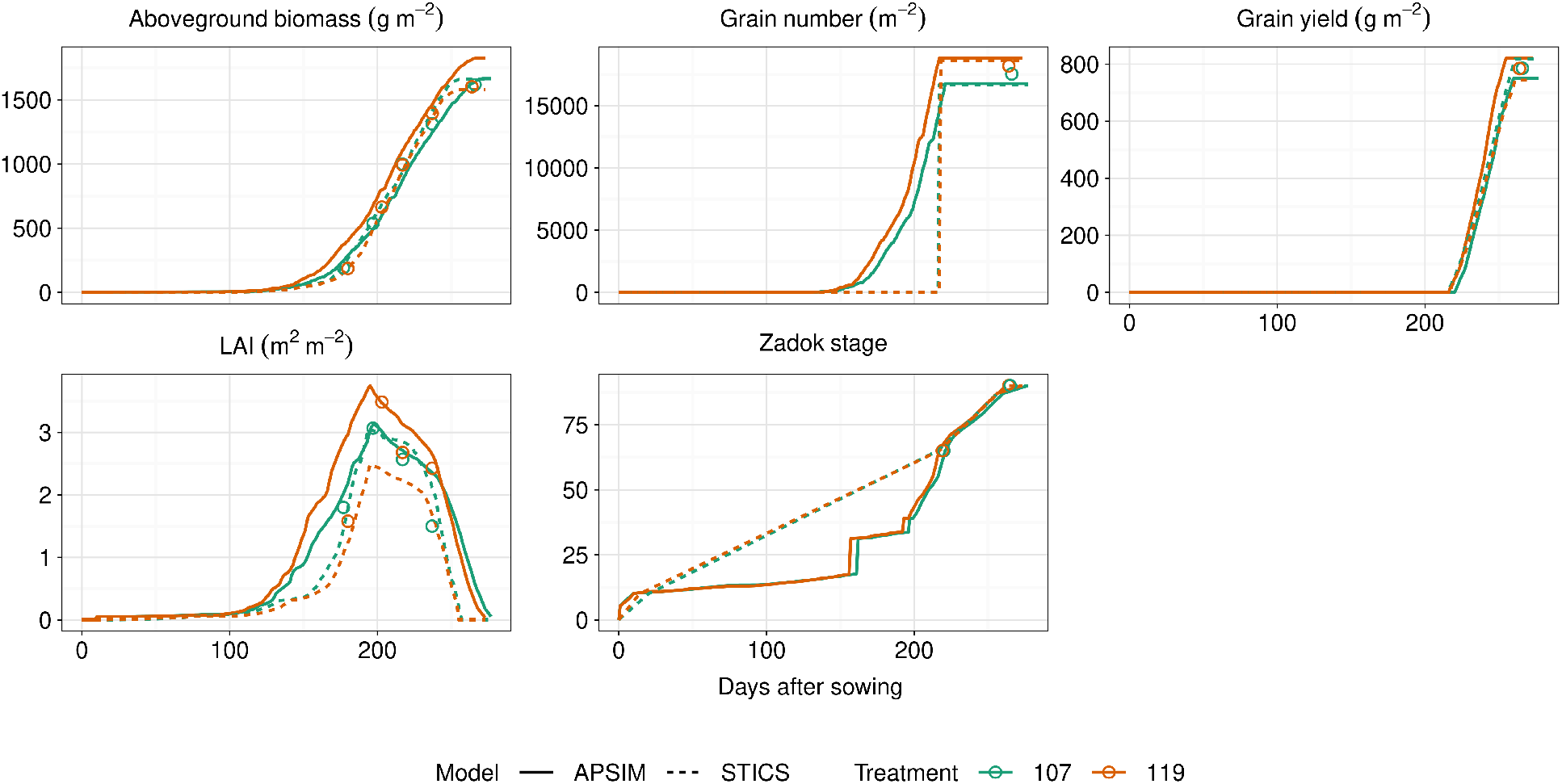
Calibration at Braunschweig using control treatments only, i.e. ambient CO_2_, no temperature increase treatment. Points indicate observation means. No measure of uncertainty was provided in this dataset.

**Figure S5:**
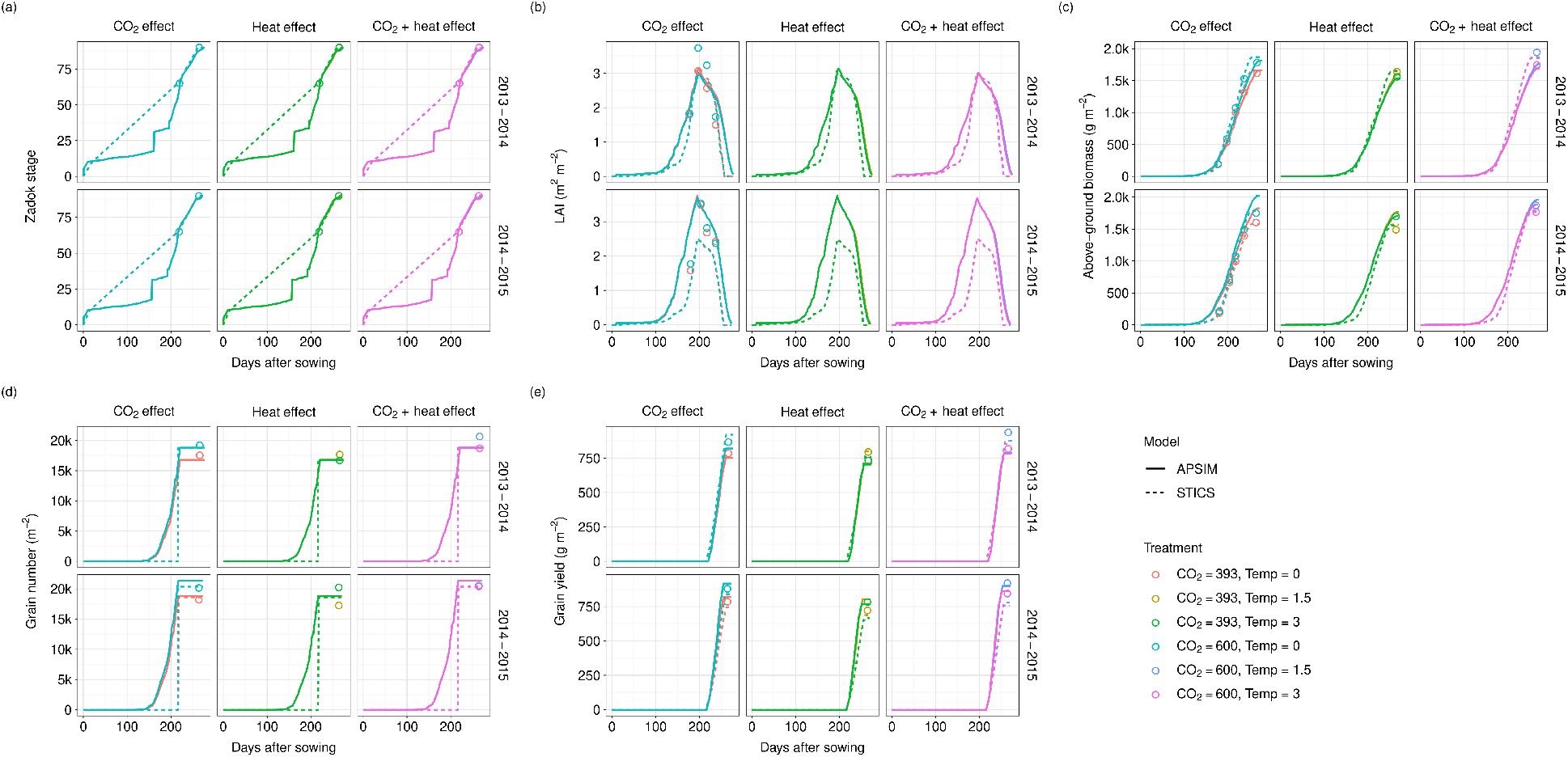
Simulated vs. observed (a) phenology (Zadok scale), (b) leaf area index (LAI), (c) above-ground biomass, (d) grain number, and (e) grain yield for all treatments in Braunschweig. Points represent observed data.

**Figure S6:**
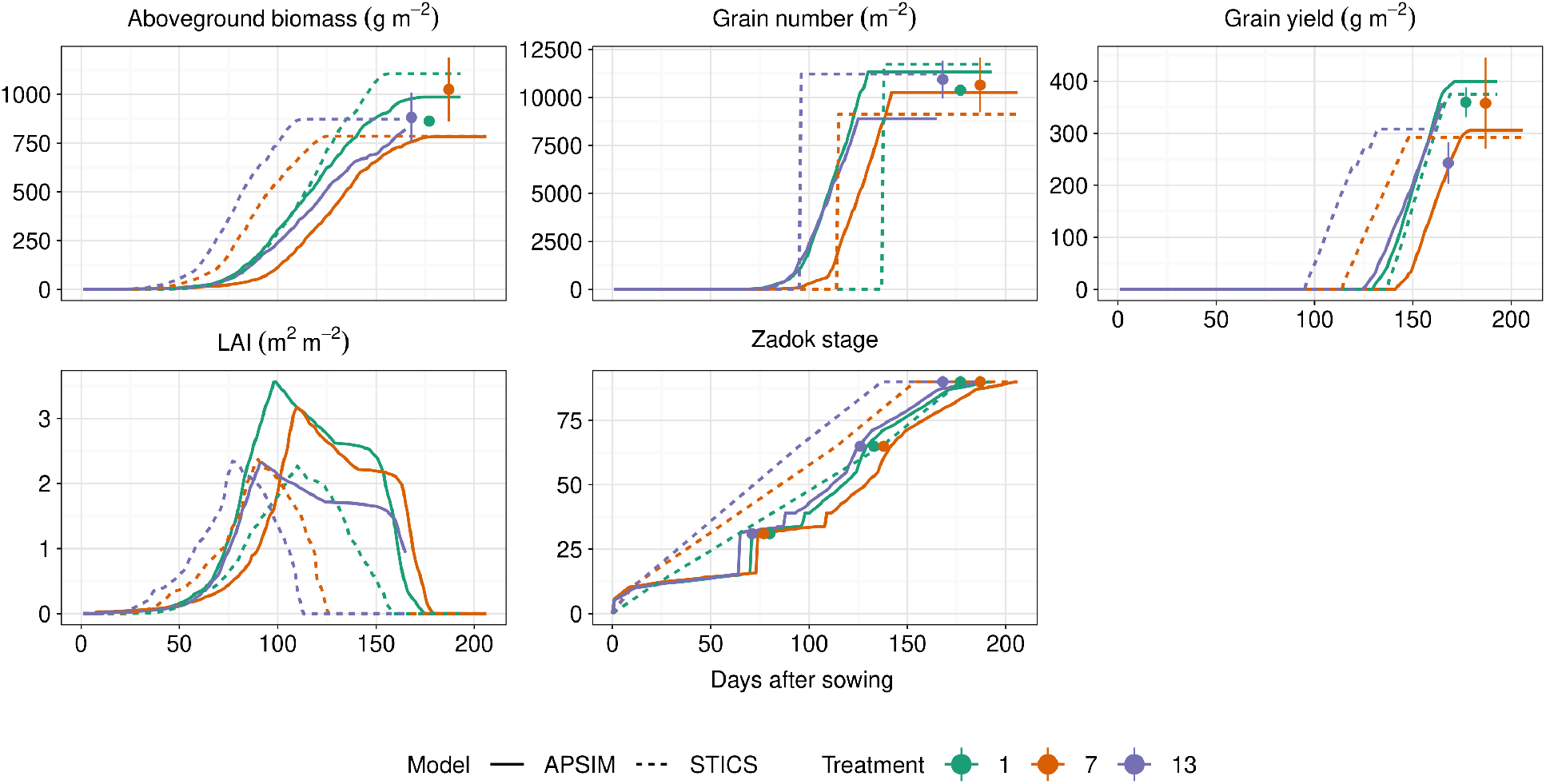
Calibration at Horsham using control treatments only, i.e. ambient CO_2_, no water stress, early sowing date. Points indicate observation means, and error bars are standard deviations.

**Figure S7:**
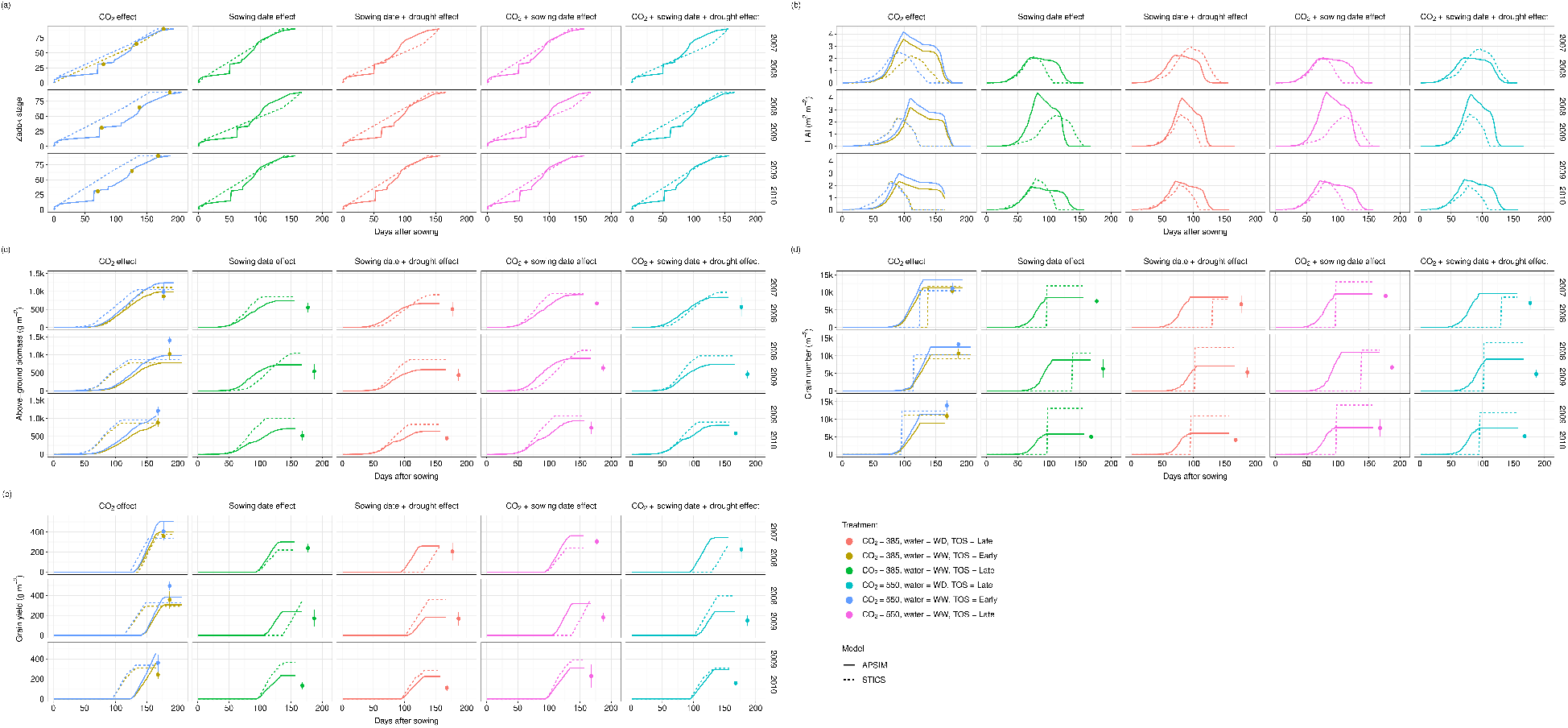
Simulated vs. observed (a) phenology (Zadok scale), (b) leaf area index (LAI), (c) above-ground biomass, (d) grain number, and (e) grain yield for all treatments in Horsham. Points represent observed data; error bars indicate standard deviations.

**Figure S8:**
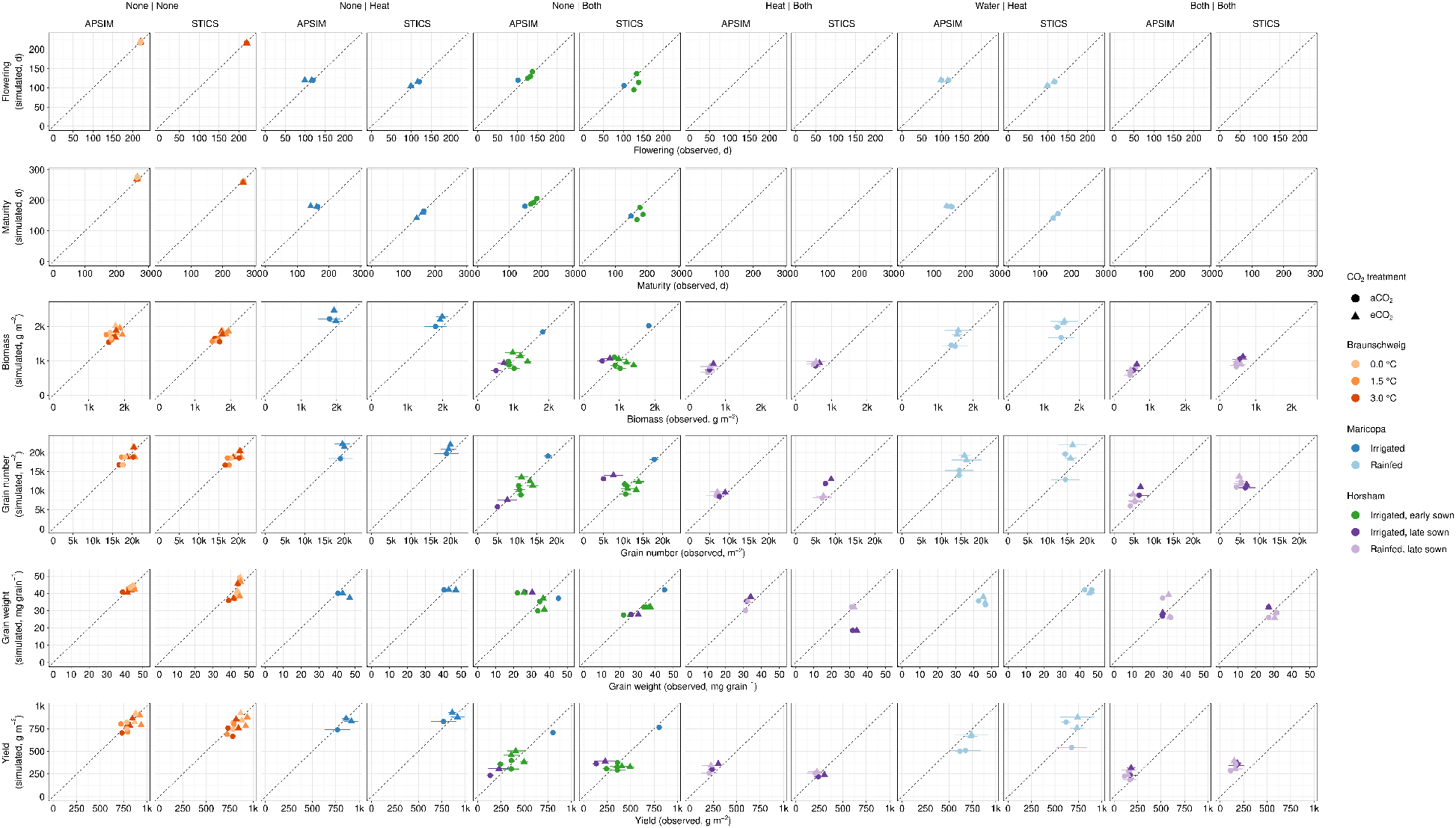
Comparison of simulated versus observed values across all variables, trials, and treatments, categorised by stress type (see Figure 8 for definitions). Point shapes distinguish CO_2_ treatments: ambient (aCO_2_, circles) and elevated (eCO_2_, triangles).

